# Cardiac pathologies in mouse loss of imprinting models are due to misexpression of H19 long noncoding RNA

**DOI:** 10.1101/2021.02.23.432554

**Authors:** Ki-Sun Park, Beenish Rahat, Zu-Xi Yu, Jacob Noeker, Apratim Mitra, Russell Knutsen, Danielle Springer, Beth Kozel, Karl Pfeifer

**Affiliations:** Division of Intramural Research, Eunice Kennedy Shriver National Institute of Child Health and Human Development, National Institutes of Health, Bethesda MD 20892 USA; Pathology Core, National Heart Lung and Blood Institute, National Institutes of Health, Bethesda, MD 20892 USA; Laboratory of Vascular and Matrix Genetics, National Heart Lung and Blood Institute, National Institutes of Health, Bethesda, MD 20892 USA; Murine Phenotyping Core, National Heart Lung and Blood Institute, National Institutes of Health, Bethesda, MD 20892 USA

**Keywords:** Beckwith Wiedemann syndrome, epigenetics, H19 lnc RNA, loss of imprinting/Igf2

## Abstract

Maternal loss of imprinting (LOI) at the *H19/IGF2* locus results in biallelic *IGF2* and reduced *H19* expression and is associated with Beckwith Wiedemann syndrome (BWS). We use mouse models for LOI to understand the relative importance of *Igf2* and *H19* mis-expression in BWS phenotypes. Here we focus on cardiovascular phenotypes and show that neonatal cardiomegaly is exclusively dependent on increased *Igf2*. Circulating IGF2 binds cardiomyocyte receptors to hyperactivate mTOR signaling, resulting in cellular hyperplasia and hypertrophy. These *Igf2*-dependent phenotypes are transient: cardiac size returns to normal once *Igf2* expression is suppressed postnatally. However, reduced *H19* expression is sufficient to cause progressive heart pathologies including fibrosis and reduced ventricular function. In the heart, *H19* expression is concentrated predominantly in endothelial cells (ECs) and regulates EC differentiation both, *in vivo* and *in vitro*. Finally, we establish novel mouse models to show that cardiac phenotypes depend on *H19* lncRNA interactions with *let7* microRNA.

## Introduction

There are100-200 imprinted genes in mammals. These genes are organized into discrete clusters where monoallelic expression is dependent upon a shared regulatory element known as the *Imprinting Control Region* (*ICR*)(Barlow & Bartolomei, 2014). Imprinted genes are frequently involved in human disease and developmental disorders (Eggermann *et al*, 2015; Feinberg & Tycko, 2004; Horsthemke, 2014; Kalish *et al*, 2014; Peters, 2014). Sometimes, these diseases are due to inactivating point mutations of the only transcriptionally active allele. Alternatively, imprinting diseases are caused by disruption of ICR function, leading to mis-expression of all genes in the cluster.

One imprinted cluster is the *IGF2/H19* locus on human chromosome 11p15.5. Imprinting in this >100 kb region is determined by the *H19ICR*, located just upstream of the *H19* promoter (Kaffer *et al*, 2000; Thorvaldsen *et al*, 1998). As described in Figure 1A, the *H19ICR* organizes the locus such that transcription of the *IGF2* (*Insulin-like <u>G</u>rowth <u>F</u>actor <u>2</u>*) and *H19* genes are expressed from the paternal and maternal chromosomes, respectively (Ideraabdullah *et al*, 2008; Murrell, 2011; Yoon *et al*, 2007). (Note that in medical genetics, the *H19ICR* is also known as Imprinting Center 1 or IC1).

**Figure 1.**
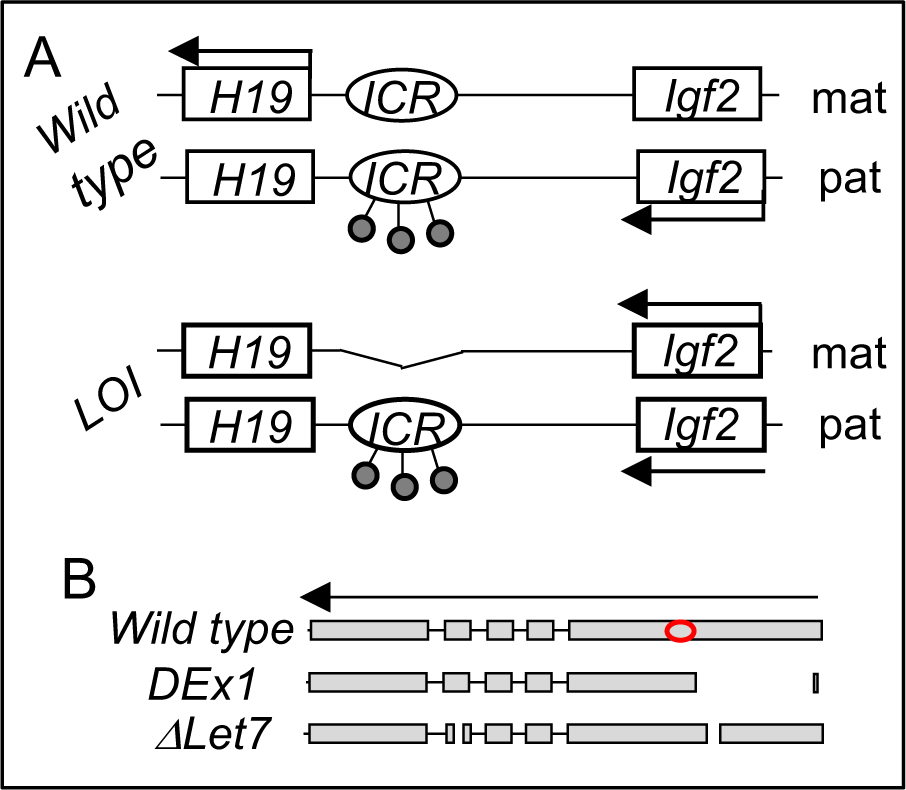
*The H19/Igf2* locus. **A**. Schematic of maternal (mat) and paternal (pat) chromosomes in wild type and in loss of imprinting (LOI) mice. Gene expression is indicated by horizontal arrows. In wild type mice, the paternal copy of the imprinting control region (ICR) is inactivated by DNA methylation (filled lollipops). **B**. Schematic of *wild type*, *ΔEx1*, and *ΔLet7 H19* alleles. *H19* exons 1-5 are shown as filled rectangles. *ΔEx1* is a 700 bp deletion at the 5’ end of exon 1. *ΔLet7* was constructed for this study by simultaneous deletion of *let7* binding sites in *H19* exons 1 and 4. The red oval identifies coding sequences for miR-675. Arrowheads show the direction of transcription.

*IGF2* encodes a peptide hormone that binds to and activates the Insulin receptor (InsR) and Insulin-like growth factor 1 receptor (Igf1R) kinases to promote cell growth and proliferation (Bergman *et al*, 2013). In contrast, the functional product of the *H19* gene is a 2.3 kb long non-coding RNA whose biochemical functions remain controversial (Brannan *et al*, 1990; Gabory *et al*, 2010). Reported roles for the *H19* lncRNA include: 1) acting as the precursor for microRNAs (miRNA-675-3p and miRNA-675-5p) (Cai & Cullen, 2007; Keniry *et al*, 2012), 2) regulating the bioavailability of *let7* miRNAs (Gao *et al*, 2014; Geng *et al*, 2018; Kallen *et al*, 2013; Li *et al*, 2015), 3) interacting with p53 protein to reduce its function (Hadji *et al*, 2016; Park *et al*, 2017; Peng *et al*, 2017; Yang *et al*, 2012; Zhang *et al*, 2019; Zhang *et al*, 2017), and 4) regulating DNA methylation to thereby modulate gene expression (Zhou *et al*, 2019; Zhou *et al*, 2015).

In humans, disruption of the maternally inherited *H19ICR* results in biallelic *IGF2* along with reduced *H19* expression and is associated with the developmental disorder, Beckwith Wiedemann syndrome (BWS) (Jacob *et al*, 2013). BWS is a fetal overgrowth disorder but the specific manifestations of overgrowth vary between patients. Cardiomegaly is a common newborn presentation but typically resolves without treatment. Cardiomyopathies are rarer and include ventricular dilation, valve/septal defects, fibrotic and rhabdomyoma tumors, and vascular abnormalities (Cohen, 2005; Descartes *et al*, 2008; Drut *et al*, 2006; Elliott *et al*, 1994; Greenwood *et al*, 1977; Knopp *et al*, 2015; Longardt *et al*, 2014; Ryan *et al*, 1989; Satge *et al*, 2005). BWS incidence correlates with artificial reproductive technologies (ART) (DeBaun *et al*, 2003; Gicquel *et al*, 2003; Halliday *et al*, 2004; Hattori *et al*, 2019; Johnson *et al*, 2018; Maher *et al*, 2003; Mussa *et al*, 2017) and among BWS patients, the frequency of heart defects is higher in those born via ART (Tenorio *et al*, 2016).

We have generated a mouse model that recapitulates the molecular loss of imprinting (LOI) phenotypes of BWS (Figure 1A) (Srivastava *et al*, 2000). That is, deletion of the *H19ICR* on the maternal chromosome results in biallelic *Igf2* and reduced levels of *H19*. In this study, we show that the LOI mouse model displays cardiovascular defects seen in BWS patients. Genetic and developmental analyses indicate that mis-expression of *Igf2* and *H19* act independently on distinct cell types to cause the cardiac phenotypes. During fetal development, increased circulating IGF2 activates AKT/mTOR pathways in cardiomyocytes resulting in cellular hypertrophy and hyperplasia. This neonatal hypertrophy is transient, non-pathologic, and unaffected by the presence or absence of a functional *H19* gene. However, loss of *H19* lncRNA results in cardiac fibrosis and hypertrophy and a progressive cardiac pathology in adult animals. In both neonatal and adult hearts, *H19* lncRNA expression is concentrated in endothelial cells (ECs). *In vivo*, loss of *H19* results in high incidence of ECs that co-express endothelial and mesenchymal markers. Similarly, primary cardiac endothelial cells can be driven toward a mesenchymal phenotype by manipulating *H19* expression levels. Thus, this research identifies a novel developmental role for the *H19* lncRNA in regulating cardiac endothelial cells. In fact, this role for *H19* in restricting endothelial cell transitions in the heart is unexpected given previous analyses of *H19* function in vitro in transformed cell lines. Finally, we describe structure-function analyses in two novel mouse models and show in that *H19* lncRNA acts by regulating *let7* bioavailability.

## Results

### Defective structure and function in hearts from mice with H19/Igf2 maternal loss of imprinting (LOI)

Wild type and LOI mice were generated by crossing *H19^ΔICR^/H19^+^* females with wild type C57Bl/6J males. (See Figure 1A for a description of the *H19ΔICR* allele). In mice (as in humans), maternal LOI results in biallelic (2X) expression of *Igf2* and reduced levels of *H19* RNA (Supplemental Figure 1A). Hearts isolated from P1 LOI mice display cellular hyperplasia and cellular hypertrophy. Hyperplasia is indicated by increased staining for Ki-67 in tissue sections (Figure 2A, C) and by increased levels of Ki-67 and of cyclins E1 and D1 in protein extracts (Figure 2D). Cellular hypertrophy is demonstrated by measuring surface areas of primary cardiomyocytes isolated from wild type and LOI neonates (Figure 2E, F). Apart from their increased size, neonatal LOI hearts do not display any obvious pathologies. For example, we did not see increased fibrosis or expression of protein markers associated with heart disease. (See Figure 3B for markers that were assayed but did not show aberrant expression). Furthermore, by 2 months of age, we were unable to distinguish LOI mice by cardiomegaly.

**Figure 2.**
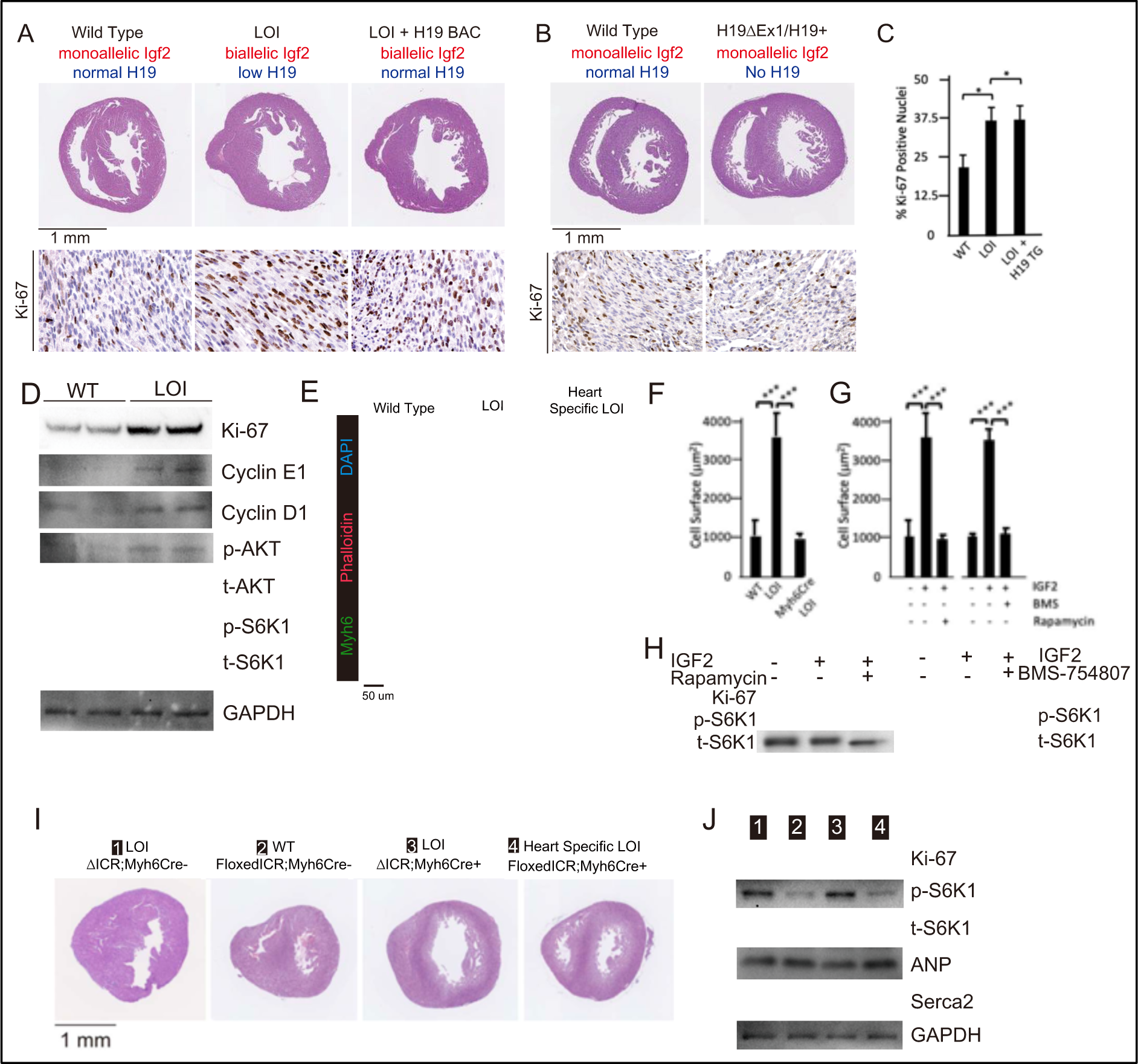
Cardiac hypertrophy in neonatal LOI mice is mediated by circulating IGF2’s activation of AKT/mTOR signaling in cardiomyocytes and is independent of *H19* gene function. **A, B.** Heart morphology in wild type, LOI, and LOI + H19 BAC littermates **(A)** or in wild type and *H19*-deficient littermates **(B)**. Top panels, transverse sections were taken from fixed hearts at 200 mm from the apex. Bottom panels, Ki-67 (brown stain) is a marker for cell proliferation. LOI, *Η19^ΔΙχρ^/H19^+^*; LOI + BAC, *H19^ΔICR^/H19^+^* mice that also carry a 140 kb Bacterial Artificial Chromosome transgene that restores normal *H19* expression. Notice the thickened walls, misshaped right ventricles, and high levels of Ki-67 expression in LOI and in LOI + BAC transgenic neonates. **C.** Quantitation of Ki-67 expression as assayed in panel A. (N = 4). **D**. Immunoblot analyses of heart extracts prepared from wild type and LOI littermates. LOI hearts show increased levels of proliferation markers, Ki-67, Cyclin EI, and Cyclin D1 and also increased levels of phosphorylated AKT and S6K1 (a target of mTORC1). **E, F**. Cardiomyocyte cellular hypertrophy in LOI animals is cell non-autonomous. Primary cardiomyocyte cultures were prepared from wild type, LOI, and from littermates carrying an *ICR* deletion only in cardiomyocytes (see below). Cells were cultured overnight, stained for MYH6 (to identify cardiomyocytes) and Phalloidin (to facilitate measurement of surface areas). For each culture (N= 5 per genotype), at least 30 cells were measured. **G, H.** Exogenous IGF2 peptide induces cellular hypertrophy in wild type cardiomyocytes through mTOR pathways. Primary cardiomyocytes were prepared from wild type neonates and cultured overnight with IGF2 before measurement of cell surface area (**G**) or preparation of protein extracts for immunoblotting (**H**). The effect of increased IGF2 is prevented by treatment with BMS 754807 or with Rapamycin. BMS inhibits IgfR1 and Ins2 receptor kinases (Carboni *et al*, 2009). Rapamcyin blocks a subset of mTOR activities (Li *et al*, 2014). **I, J.** LOI phenotypes in cardiomyocytes are cell non-autonomous. *H19^ICRflox^/H19^DICR^* females were crossed with males carrying the *Myh6Cre* transgene to generate 4 kind of pups: *H19^ΔICR^/H19^+^* (#1) and *H19^ΔICR^/H19^+^ Myh6Cre* (#3) will display LOI in all cell types; *H19^ICRflox^/H19^+^* (#2) will display wild type expression patterns for *Igf2* and *H19*; and *H19^ICRflox^/H19^+^ Myh6Cre* mice will show LOI only in cardiomyocytes. Hearts were analyzed for cellular hypertrophy (**E)**, megacardia and hyperplasia (**I**), and protein expression (**J**). In all assays, *H19^ICRflox^/H19^+^ Myh6Cre* mice were indistinguishable from their wild type littermates. All bar graphs show mean ± SEM. *, p<0.05; ***, p<0.001 (Student’s t-tes). LOI, Loss of imprinting (*H19^ΔICR^/H19^+^*).

**Figure 3.**
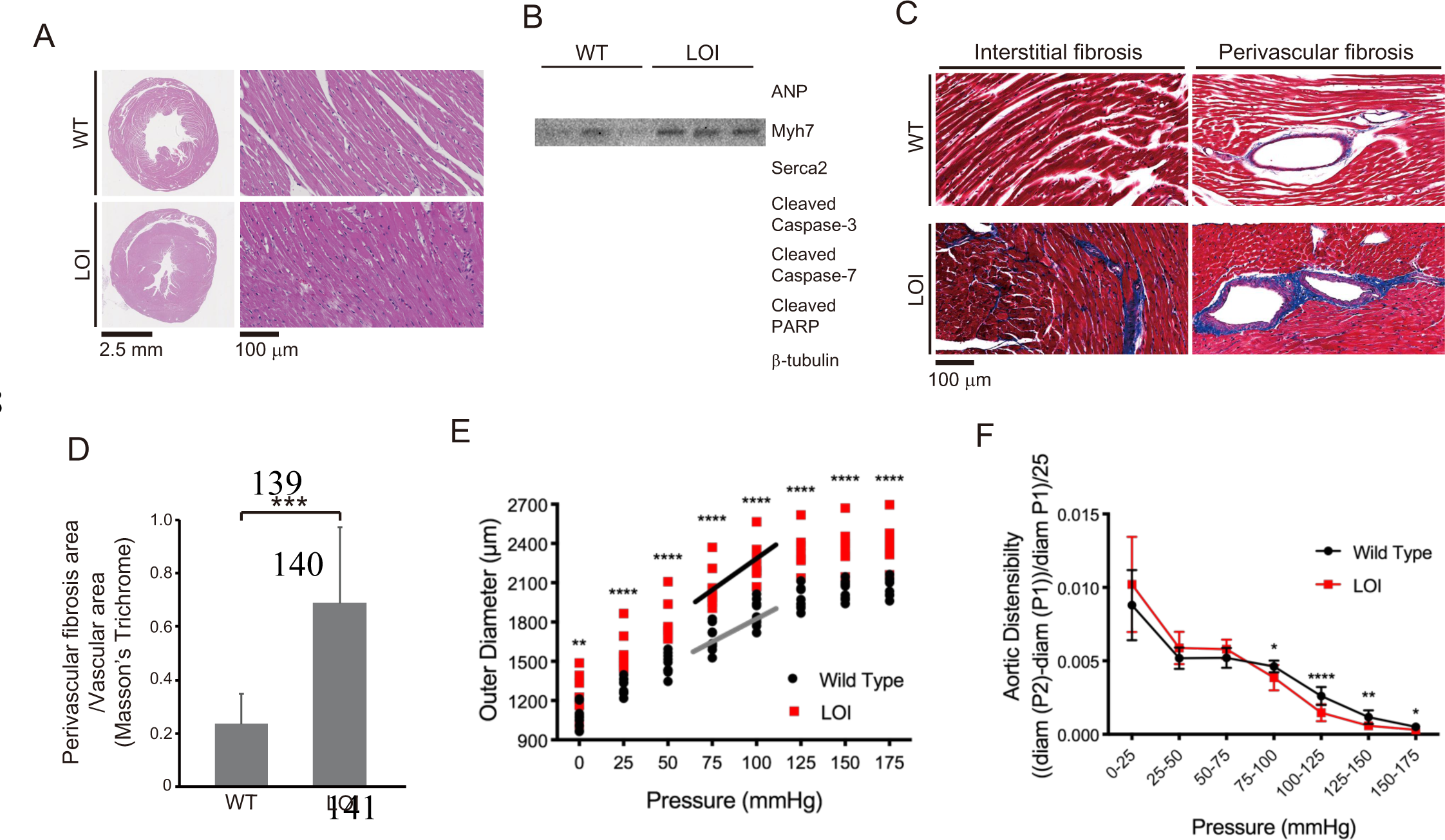
Cardiomyopathies in adult LOI mice. **A.** Transverse sections were collected midway along the longitudinal axis from hearts collected from 6-month-old wild type (WT) and LOI mice and stained with hematoxylin and eosin. **B.** Immunoblot analyses of whole heart extracts prepared from 1-yeart WT and LOI mice. Note the altered expression of ANP (Atrial Natriuretic Peptide), Myh7 (Myosin Heavy Chain 7), Serca2 (Sarco/endoplasmic reticulum Ca^++^ ATPase), Cleaved Caspase-3, and Cleaved Poly ADP Ribose Polymerase (PARP). b-tubulin is a loading control. **C, D**. Masson’s trichrome staining of sections described in panel **A**. Red, muscle fibers; blue, collagen. Sections from 5 wild type and 5 LOI animals were used to calculate fibrosis. Bar graphs show mean ± SEM. Data were analyzed by Student’s t-test. **E, F.** Ascending aortas were isolated from 10 wild type and 8 LOI mice and pressure-diameter curves generated. **E.** Increased diameters across a wide range of applied pressures. **F.** Increased segmental distensibility across physiologically relevant pressures. Data were analyzed by two-way repeated measure ANOVA. For all panels: *, P<0.05; **, P<0.01; ***, P<0.001; ****, P<0.0001.

We continued to monitor cardiovascular phenotypes in LOI and wild type mice until 19 months of age. By 6 months, LOI mice displayed cardiac hypertrophy as measured by a 28% increase in heart weight/tibia length ratios (wild type = 10.0 ± 1.7 mg/mm, N= 8; LOI = 12.8 ± mg/mm, N=10; p = 0.005). Transverse sections revealed increased fiber diameter in LOI hearts (wild type = 10.2 ± 0.7 μm; LOI = 14.4 ± 0.8 μm; p = 0.007) (Figure 3A). Cardiac hypertrophy is often a poor prognostic sign and is associated with most forms of heart failure (Heinzel *et al*, 2015; Vakili *et al*, 2001). However, hypertrophy can also be physiologic (McMullen & Jennings, 2007; Shimizu & Minamino, 2016). The hypertrophy in LOI mice might be considered pathologic based on increased levels of ANP, Myh7, cleaved Caspace-3, cleaved Caspace-7, and cleaved PARP proteins as well as decreased levels of Serca2 protein in all LOI mice by 1 year of age (Figure 3B) (Mitra *et al*, 2013; van Empel *et al*, 2005). Finally, both interstitial and perivascular fibrosis are prominent in LOI animals by 6 months of age (Figure 3C, D).

Table 1 summarizes echocardiography phenotypes from 13-month-old mice. Left ventricles (LV) from LOI mice are dilated (as measured by increased LV volumes at both systole and diastole), mildly hypertrophic (as measured by increased wall thickness, LVAW diastole and LVPW diastole), and show diminished function (as measured by reduced ejection fractions, % EF). LOI mice showed large increases in velocity and turbulence of blood flow from the LV outflow tract. Finally, major vessel lumen diameters (measured at the aortic arch and the first brachial arch) were >30% larger in LOI mice.

**Table 1.** Echocardiography of wild type (WT) and loss of imprinting (LOI) mice at 13 and at 16 months. LV, left ventricle; EF, ejection fraction; AW, anterior wall; PW, posterior wall; OT, outflow tract. P-value is by Student’s t-test.

| Phenotype | 13 months |  |  |  | 16 months |  |  |  |
| --- | --- | --- | --- | --- | --- | --- | --- | --- |
| | Mean ± SEM | | P-value | % Change | Mean ± SEM | | P-value | % Change |
|  | WT (N = 11) | LOI (N = 10) |  |  | WT (N = 11) | LOI (N = 9) |  |  |
| Heart rate (bpm) | 504 ± 12 | 499 ± 11 | 0.77 | -1 | 529 ± 20 | 540 ± 16 | 0.77 | 2 |
| LV Volume Systole (μl) | 24.1 ± 1.5 | 39.2 ± 4.7 | 0.01 | 63 | 28.3 ± 1.4 | 46.6 ± 1.6 | <0.001 | 65 |
| LV Volume Diastole (μl) | 67.9 ± 3.0 | 79.8 ± 4.6 | 0.05 | 17 | 74.2 ± 2.8 | 91.3 ± 3.3 | <0.001 | 23 |
| LV EF (%) | 64.7 ± 1.2 | 51.9 ± 3.8 | <0.01 | -19 | 62.5 ± 0.9 | 48.6 ± 1.7 | <0.001 | -22 |
| LVAW Systole (mm) | 1.40 ± 0.01 | 1.44 ± 0.02 | 0.09 | 3 | 1.39 ± 0.01 | 1.52 ± 0.04 | 0.01 | 8 |
| LVAW Diastole (mm) | 0.88 ± 0.01 | 1.01 ± 0.03 | <0.01 | 15 | 0.91 ± 0.01 | 1.08 ± 0.04 | <0.001 | 20 |
| LVPW Systole (mm) | 1.35 ± 0.01 | 1.39 ± 0.02 | 0.17 | 3 | 1.34 ± 0.01 | 1.42 ± 0.03 | 0.4 | 6 |
| LVPW Diastole (mm) | 0.87 ± 0.02 | 0.98 ± 0.04 | 0.02 | 12 | 0.88 ± 0.02 | 1.08 ± 0.04 | <0.001 | 19 |
| LVOT mean gradient | 2.4 ± 0.2 | 6.8 ± 1.8 | 0.04 | 185 | 2.3 ± 0.2 | 4.9 ± 1.1 | 0.04 | 115 |
| LOVT mean velocity | 768 ± 30 | 1211 ± 162 | 0.02 | 58 | 750 ± 37 | 1066 ± 112 | 0.02 | 42 |
| LVOT peak gradient | 5.8 ± 0.3 | 15.7 ± 3.7 | 0.03 | 170 | 5.5 ± 0.5 | 13.3 ± 3.3 | 0.04 | 143 |
| LVOT peak velocity | 1201 ± 35 | 1870 ± 215 | 0.01 | 56 | 1158 ± 52 | 1732 ± 200 | 0.02 | 50 |
| Aorta Systole (mm) | 1.65 ± 0.04 | 2.14 ± 0.10 | <0.01 | 30 | 1.70 ± 0.04 | 2.25 ± 0.12 | 0.002 | 32 |
| Aorta Diastole (mm) | 1.44 ± 0.04 | 1.86 ± 0.07 | <0.01 | 30 | 1.48 ± 0.04 | 2.07 ± 0.13 | 0.002 | 40 |
| First Brachial Arch (mm) | 0.78 ± 0.03 | 1.06 ± 0.07 | <0.01 | 36 | 0.78 ± 0.02 | 1.09 ± 0.08 | 0.004 | 40 |

Scatterplots of echocardiography data from 13 month animals show that most LOI phenotypes are heterogeneous and are not normally distributed (Supplemental Figure 2). Rather phenotypes for volume, mass, ejection fraction, and outflow tract velocity and turbulence are all bimodal: 6-7 animals display mild phenotypes and 3-4 animals display extreme pathologies (Supplemental Figure 2A-D). The only exception to this pattern is seen in arterial diameter phenotypes. In this case, the variance among LOI animals is low (like their WT cohorts) and all the LOI animals display a pathologic phenotype (Supplemental Figure 2E, F).

Supplemental Table 1 summarizes correlations between the various phenotypes identified by echocardiography. Cardiac function as measured by ejection fraction is inversely correlated with LV volume (RR = 0.81). However, function correlates only moderately with wall thickness (RR = 0.53) and not at all with outflow tract defects (RR = 0.02), or with aortic diameter (RR <0.01). Thus, LOI associated phenotypes are not uniformly penetrant. Rather, each mouse presents a distinct array of defects. The only invariant is that all LOI mice have arterial diameters larger than their wild type counterparts.

The right-hand columns in Table 1 summarize echocardiography results from the same mice at 16 months of age. We observed the same ventricular abnormalities: reduced ejection fraction, increased chamber size, and increased wall thickness. However, on scatter plots we see that wild type and mutant animals now show non-overlapping phenotypes, consistent with the idea that ventricular failure is progressing in LOI mice (Supplemental Figure 2). Note that the LOI mouse with the poorest function at 13 months (25 % EF) died prior to this second analysis.

Finally, in vivo analyses at 19 months identified significant pathological reductions in both systolic blood pressure (WT = 105 ± 2, LOI = 93 ± 3, p = 0.01) and pulse pressure (WT = 37 ± 1, LOI = 28 ± 1, p <0.001) in mutant mice (Supplemental Figure 3). These data confirm that *H19/Igf2* LOI has a substantial effect on cardiovascular function.

As described above, increased artery diameter is a phenotype where by 13 months, LOI and WT mice sorted into phenotypically distinct cohorts. This suggested that abnormal blood vessel structure might be a relatively primary defect. We focused additional attention to this phenotype and measured outer diameters of isolated ascending aorta and carotid arteries in response to applied pressures on a pressure myograph (Figure 3E, Supplemental Figure 4A). Arteries from LOI mice are larger in diameter across all applied pressures. Moreover, across normal physiological pressure ranges (75-125 mmHg) arteries from mutant mice are more sensitive to changes in pressure and lumens reach their maximum diameter at lower pressures. They are appropriately distensible at low (elastic) pressures but are stiffer than WT vessels over higher pressure intervals, including most physiologic pressures (Figure 3F, Supplemental Figure 4B).

In sum, *H19/Igf2* LOI in mice results in transient neonatal cardiomegaly and then a progressive cardiomyopathy. Note that results shown in Figure 3 and in Table 1 describe comparisons of age-matched male mice. Adult LOI females consistently showed relatively weak phenotypes and p values were not significant (data now shown). However, neonatal hypertrophy and hyperplasia occurs in both male and female pups. This apparent paradox was the first clue that the relationship between the neonatal hypertrophy and the adult disease phenotypes was not straightforward.

### Hypertrophy and hyperplasia in neonatal LOI mice is dependent on hyperactivation of mTOR/AKT signaling by increased dosage of IGF2 peptide

To understand the specific roles for mis-expression of *Igf2* and of *H19* in neonatal cardiomegaly we performed two genetic analyses. First, we rescued *H19* expression in an LOI background by introducing a 140 kb H19 Bacterial Artificial Chromosome (H19 BAC) (Kaffer *et al*, 2001; Kaffer *et al*., 2000) but still saw cardiomyocyte hypertrophy and hyperplasia in neonates (Figure 2A, C). Second, we tested the effect of removing *H19* in a background where *Igf2* remains monoallelic by comparing *H19^ΔEx1^/H19^+^* pups (Figure 1B) with wild type littermates. Loss of *H19* lncRNA does not result in neonatal cardiomyocyte hypertrophy or hyperplasia (Figure 2B). Altogether, we conclude that loss of *H19* lncRNA does not contribute to neonatal hypergrowth. Rather, this neonatal hypertrophy is dependent only upon biallelic (2X dosage) *Igf2* transcription.

IGF2 peptide works by binding and activating InsR and IgfR kinases and mTOR/AKT signaling is a known downstream target of these receptor kinases (Bergman *et al*., 2013). In addition, studies document the role of AKT/mTOR signaling in cardiomyocyte cell division and hypertrophy (Sciarretta *et al*, 2014). Consistent with a critical role for AKT/mTOR signaling in LOI-dependent neonatal hypertrophy, hearts from LOI neonates show increased levels of phosphorylated AKT and of phosphorylated S6K1, a downstream marker for mTORC1 activity (Figure 2D). Moreover, the LOI cellular hypertrophy and pAKT hyperactivation phenotypes can be phenocopied by treatment of wild type primary cardiomyocytes with IGF2 peptide. However, IGF2 action is blocked by BMS-754807, a specific inhibitor of the receptor kinase, or by treatment with rapamycin, an mTOR signaling pathway inhibitor (Figure 3G, H).

*Igf2* is widely expressed in the embryo. In fact, expression of *Igf2* is low in the heart relative to other tissues, especially liver and skeletal muscle (Supplemental Figure 1B). To assess the role of biallelic *Igf2* in the cardiomyocytes themselves, we crossed *H19^ΔICR^/H19^ICRflox^* females with *H19^+^/H19^+^* males carrying the Myh6Cre transgene. *H19^ICRflox^* is an allele where the *H19ICR* is flanked with *loxP* sites so that cre recombination results in deletion of the *ICR* (Srivastava *et al*., 2000). We used PCR analyses to demonstrate that the Myh6Cre transgene drives efficient ICR deletion in the heart but not in other tissues tested (skeletal muscle, liver, kidney, brain, thymus, spleen, and lung). Our cross generated wild type mice (*H19^ICRflox^/H19^+^*) and two kinds of LOI controls (*H19^ΔICR^/H19^+^; +Myh6Cre* and *H19^ΔICR^/H19^+^*) that we compared with experimental mice that had cardiomyocyte specific LOI (*H19^ICRflox^/H19^+^;+Myh6Cre*). Cardiomyocyte specific ICR deletion does not cause hypertrophy. Rather, *H19^ICRFlox^/H19^+^ Myh6Cre* mice were indistinguishable from their wild type littermates (Figure 2E, F, I, J).

### Cardiac disease in adults is dependent only upon loss of H19 lncRNA expression. Biallelic Igf2 and the resultant hypertrophy in neonatal hearts are not relevant to the adult LOI phenotype

While the *H19* BAC transgene does not prevent neonatal cardiomegaly, it does successfully prevent adult pathologies. That is, hearts from 6-month LOI mice carrying the *H19* transgene are not enlarged as determined by heart weight/tibia length ratios (LOI = 12.9±0.6 mg/mm, N=3; LOI + H19 BAC Transgene = 11.5±0.7, N=3; p <0.05), are not fibrotic (Figure 4A, B), and do not express cardiomyopathy markers (Figure 4C). Thus, loss of *H19* is <u>necessary</u> to induce LOI cardiomyopathies.

**Figure 4.**
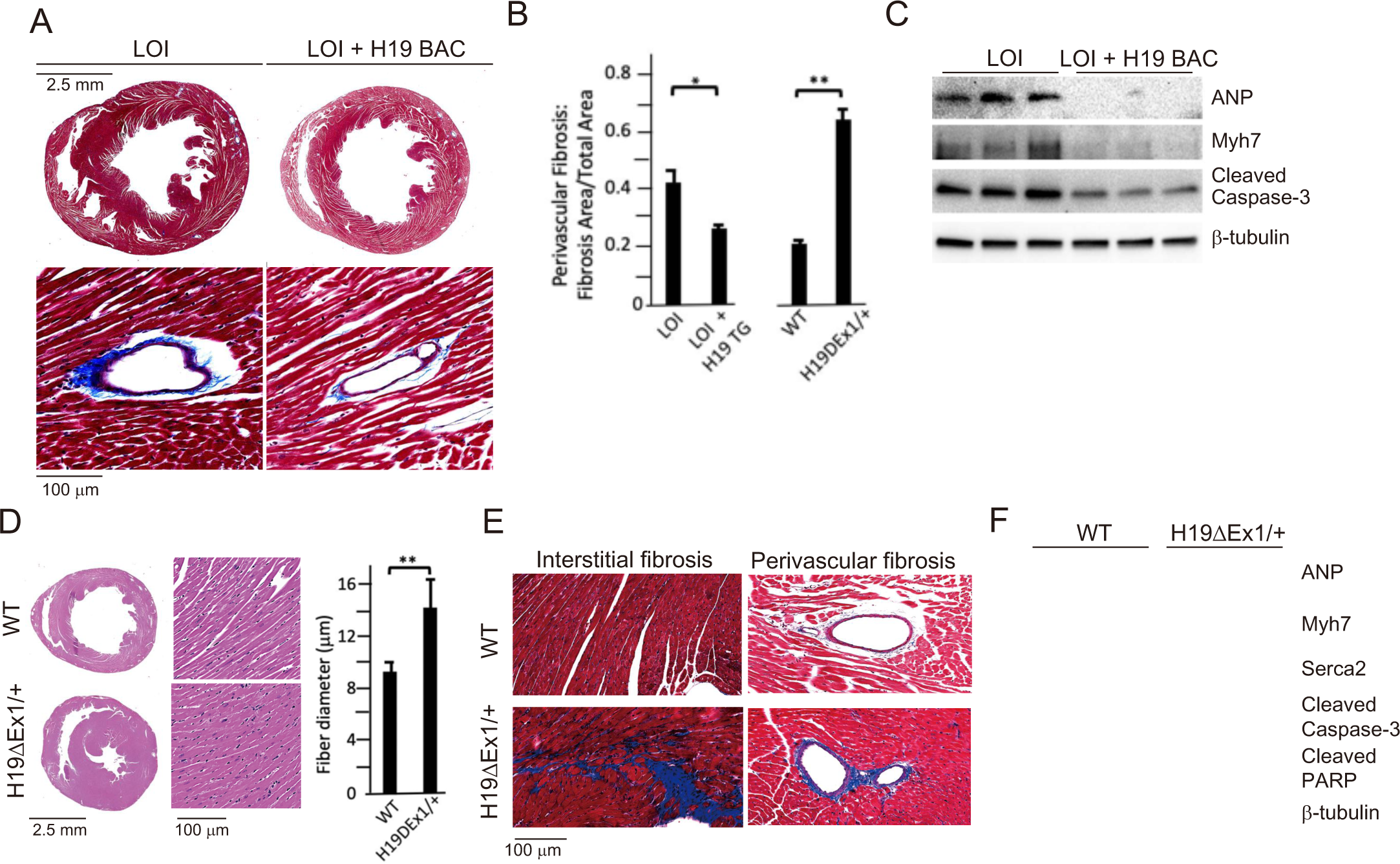
LOI pathologies in adult mice are *H19*-dependent. **A, B, C** An *H19* transgene rescues pathologies in LOI mice. Phenotypes of LOI mice or their LOI littermates that also carry an H19 Bacterial Artificial Chromosome transgene that restores wild type levels of *H19* RNA (LOI + H19 BAC). **D, E, F** *H19* deletion is sufficient to cause cardiac pathologies. Phenotypes in wild type (WT) mice and in littermates carrying the *H19ΔEx1* deletion. For histology (**A, B, D, E**) hearts were isolated from 6-month old animals and transverse sections collected midway along the longitudinal axes before staining with hematoxylin and eosin (**D**) or with Masson’s trichrome **(A, B, E)**. Bar graphs show mean + SEM. *, P<0.05; **, P<0.01 (Student’s t-test). For immunoblotting (**C, F)**, hearts were isolated from 1-year animals and investigated for ANP, Myh7, Serca2, Cleaved Caspase-3, and Cleaved PARP. b-tubulin is a loading control.

We next investigated whether loss of *H19* is <u>sufficient</u> to induce pathologies and also investigated exactly which *H19* RNA was important. The *H19* gene encodes a 2.3 kb lncRNA which is exported to the cytoplasm but also is the precursor for microRNAs, *miR-675-5p* and *miR-675-3p* (Cai & Cullen, 2007). Since LOI mice show reduced levels of both the lncRNA and of *miR-675* and because the H19 BAC transgene restores expression of both the lncRNA and the *miR-675* microRNAs, these models were not helpful in determining which RNA species prevents cardiac pathology. The *H19ΔEx1* allele is a 700 bp deletion of the 5’ end of exon 1 that leaves bases encoding the *miR-675* intact (Figure 1B). This *ΔEx1* deletion does not prevent *H19* transcription but rather, reduces *H19* lncRNA levels by destabilizing the truncated transcript (Srivastava *et al*, 2003), raising the possibility that the *ΔEx1* mutation might affect only the lncRNA. In fact, we show here that levels of *miR-675-5p* and *-3p* are unaltered in *H19^ΔEx1^/H19^+^* hearts (Supplemental Figure 5). Yet, 6 month old *H19^ΔEx1^/H19^+^* male mice display LOI cardiac pathologies including hypertrophy (Figure 4D), fibrosis (Figure 4B, E), and expression of disease markers (Figure 4F). Thus, we conclude that loss of *H19* lncRNA is sufficient to induce cardiomyopathy in adult mice.

In addition to establishing the critical importance of *H19* lncRNA, these genetic experiments also uncouple neonatal hypertrophy and adult pathology: neonatal LOI + H19 BAC mice show hypertrophy but do not develop adult pathologies while neonatal *H19^ΔEx1^/H19^+^* mice have normal sized hearts but do develop pathologies. Thus, neonatal cardiomegaly is not a risk factor for adult pathologies.

### H19 lncRNA regulates the frequency of endothelial to mesenchymal transition in mice and in isolated primary endothelial cell cultures

*H19* expression is not uniform throughout the heart but rather restricted to endothelial cells (ECs) (Figure 5A, 5B). In fetal and neonatal hearts *H19* is expressed in all endothelial cells including microvasculature. In adults, *H19* expression is restricted to endocardium and endothelial cells lining major coronary vessels (Figure 5A) (Supplemental Figure 1B). Localization was confirmed in vasculature by co-staining for both endothelial and smooth muscle markers. For example, in coronary vessels, *H19* RNA expression exclusively overlapped with endothelial specific marker von-Willebrand’s factor (Figure 5B).

**Figure 5.**
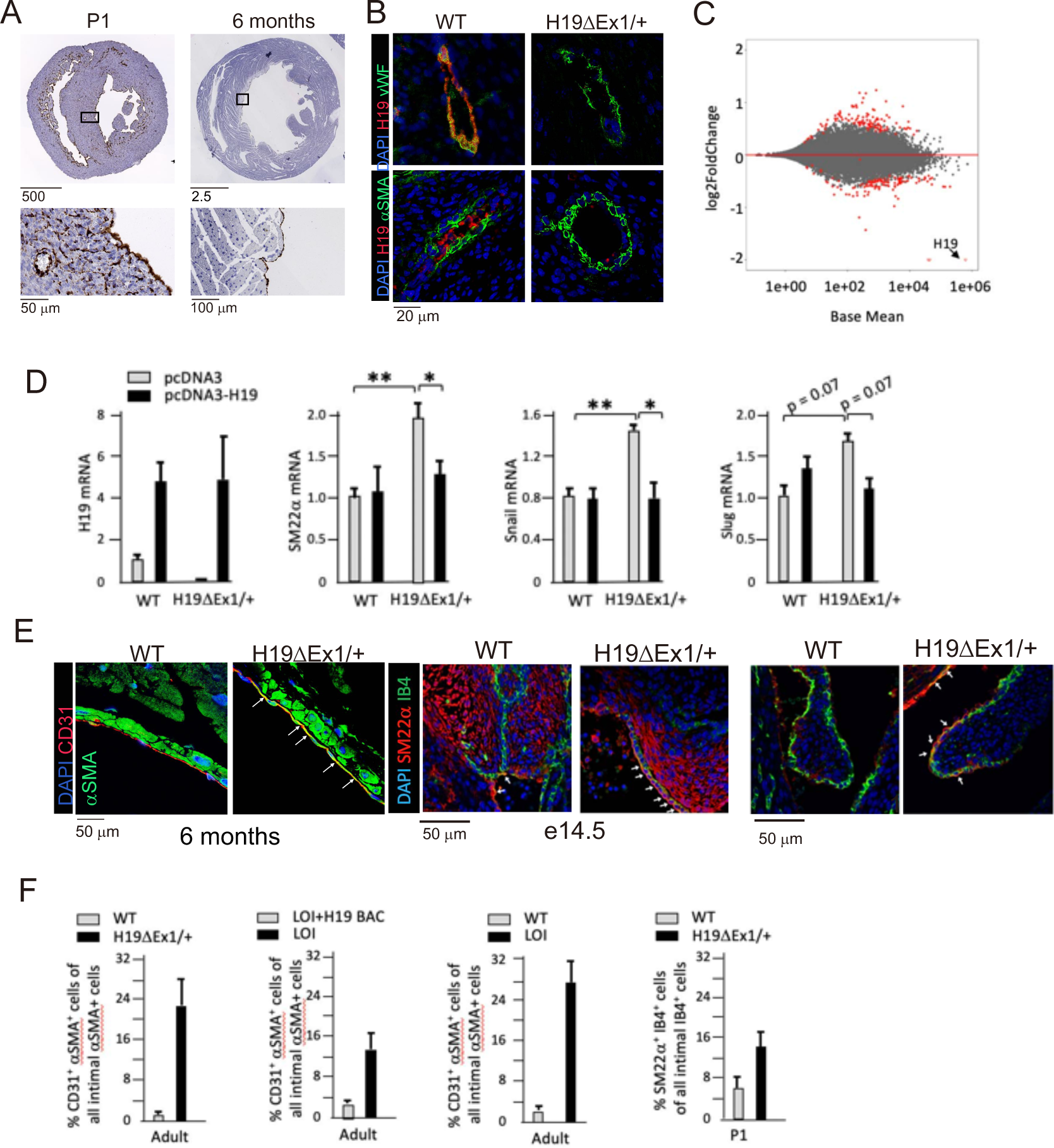
*H19* influences gene expression and cell fate in cardiac endothelial cells. **A.** In situ staining for *H19* (brown) in hearts from wild type P1 neonates or 6-month adults. **B.** Combined in situ and immunohistochemistry for hearts isolated from wild type and *H19ΔEx1/H19+* littermates at 6 months shows expression of *H19* is concentrated in endothelial cells. Sections were stained for *H19* lncRNA and then with antibodies to the endothelial marker, vWF (von Willebrand Factor), or to the smooth muscle marker, a-SMA (alpha smooth muscle actin). **C.** MA blot showing differences in expression of polyadenylated RNAs in *H19*-deficient endothelial cells. Endothelial cells were isolated from wild type (N = 4) and *H19^ΔEx1^/H19^+^* (N = 3) P2 neonatal hearts based on CD31 expression. RNAs were isolated and polyadenylated transcripts were quantitated. Genes marked in red are significantly differentially expressed at FDR<0.05. **D.** Transient transfection of *H19^ΔEx1^/H19^+^* cardiac endothelial cells with an *H19*-expression vector rescues expression of key EndMT genes. Cardiac endothelial cells were isolated from wild type and *H19*-deficient P2 hearts as described in **D** and transfected with empty expression vector (pcDNA3) or with pcDNA3 carrying mouse *H19* gDNA (pcDNA3-H19). After 24 hours in culture, RNA was extracted and cDNAs synthesized and analyzed for *H19*, *SM22a*, *Snail*, or *Slug*. For each gene, cDNA levels were normalized to *GAPDH* and then to the levels seen in wild type cells transfected with pcDNA3 only. **E, F**. Increased frequency of EndMT transitioning cells in *H19*-deficient mice. **E.** Hearts from wild type and *H19^ΔEx1^/H19^+^* littermates were isolated at e14.5, P1, and at 6 months. Sections were probed for endothelial cell markers (CD31 or IB4) and for mesenchymal markers (aSMA or SM22a) to identify cells co-expressing these genes. **F.** Frequencies of cells co-expressing endothelial and mesenchymal markers in adult and P1 hearts. The role of *H19* was determined by three independent comparisons: wild type vs. *H19^ΔEx1^/H19^+^*, LOI vs LOI + H19 BAC, wild type vs LOI. **D, F**, means ± SEM. are depicted. *, P<0.5; **, P<0.01, ***, P<0.001 (Student’s t-test).

To identify a possible function for *H19* RNA, we performed transcriptomic analyses comparing RNAs isolated from wild type and *H19*-deficient P1 hearts. Using whole heart extracts, we did not identify significant differences in gene expression. We next compared RNAs isolated from purified ECs. Hearts were dissociated into single cells using enzyme digestion and mechanical agitation and then endothelial cells were isolated based on expression of CD31 antigen. About 30,000 cells per neonatal heart were isolated to >95% purity. RNA sequencing identified 228 differentially expressed genes (DEGs) with adjusted p values of <0.1, including 111 upregulated and 117 downregulated transcripts (Figure 5C). GO analysis for biological, cellular, and molecular pathways give evidence for a change in cellular identity (Supplemental Figure 6). Specifically, enriched biological pathways included positive regulation of mesenchymal cell proliferation, positive regulation of endothelial cell migration, and cell adhesion (n = 36, p-adj = 0.003). Cellular pathways showed enrichment for genes coding for extracellular matrix (n = 42, p-adj = 1.83E-10). Enriched molecular function categories include extracellular matrix binding and TGF*β* binding (n = 7, p-adj = 0.0001; n = 5, p-adj = 0.1), as well as other pathways that are especially active during endothelial to mesenchymal transition (EndMT). EndMT is not an identifiable GO term, however, we conducted a PubMed search of the 188 DEGs described in the PubMed literature database and noted that 63 DEGs were implicated in EndMT as either players in driving the transition or as markers. Some examples include *Transforming growth factor beta receptor 3 (Tgfbr3,* up 1.5X, padj = 0.01*), Collagen Type XIII α1 chain* (*Col13a1,* up 2.0X, padj = 4.2E-09), *Bone Morphogenic Protein 6* (*bmp6*, down 0.6X, padj = 7.0E-05), *Latent Transforming Growth Factor Binding Protein 4 (Ltbp4, down 0.5x, padj = 0.008), Connective Tissue Growth Factor* (*Ctgf*, down 0.7X, padj = 0.06), *Slit Guidance Ligand 2, (Slit2,* up 1.6X, padj = 2.5E-05, *α*2 macroglobin (*α2m*, down 0.6X, 5.6E-05). Due to the results of the GO term analysis as well as the PubMed search, we speculated that *H19* might play a role in regulating EndMT.

To directly test the role of *H19* in regulating EC gene biology, we isolated primary ECs from wild type and *H19*-deficient P2 littermates and transfected with an *H19* expression vector or with an empty control vector and then assayed gene expression after 24 hours. *H19* expression reduces expression of a mesenchymal cell marker (*SM22α*) and of genes encoding transcription factors critical for EndMT (*Snail* and *Slug*) (Figure 5D).

EndMT is an essential part of the normal development of many tissues/organs including heart. For example, EndMT is critical in cardiac valve development (Kisanuki *et al*, 2001; Markwald *et al*, 1977). Studies also report that EndMT contributes to cardiac diseases including cardiac fibrosis, valve calcification, and endocardial elastofibrosis (Evrard *et al*, 2016; Goumans *et al*, 2008; Piera-Velazquez *et al*, 2011; Zeisberg *et al*, 2007). During the actual EC transition, cells will transiently express endothelial markers (like CD31or IB4) simultaneously with mesenchymal markers (like aSMA or SM22a). To understand the impact of *H19*-deficiency on EC transition *in vivo* we fixed and sectioned hearts isolated at several developmental stages from *H19^ΔEx1^/H19^+^* mice and their wild type littermates mice and looked for co-staining of these endothelial and mesenchymal markers. At each stage, we focused on the regions of the heart where *H19* expressing cells were particularly abundant, assuming that this is where a phenotype would be most readily observed. In e14.5 embryos we looked at endocardium, epicardium, valves, and blood vessels. In P1 embryos we looked at endocardium, valves, and blood vessels. In adult hearts we looked at endocardium. Comparable sections for wild type and mutant mice were identified by a cardiac pathologist blinded to genotype before we stained for EC and mesenchymal markers. In each stage we noted significant changes in co-staining frequency indicating that the likelihood of EC cell transition is increased in the absence of *H19* (Figure E, F). We confirmed these results through independent analyses that compared LOI mice with their wild type littermates (*H19^ΔICR^/H19+* vs. *H19^+^/H19^+^*) and that compared LOI mice with LOI littermates that also carried the H19 BAC transgene (*H19^ΔICR^/H19^+^* vs *H19^ΔICR^/H19^+^* + H19 BAC Transgene (Figure 5F).

### Let-7 binding sites on the H19 lncRNA are essential for normal cardiac physiology

*H19* lncRNA is known to physically bind *let-7g* and *-7i* in exon 1 and *let-7e* in exon 4 in mice (Kallen *et al*., 2013). One proposed mechanism for *H19* lncRNA function is that it regulates *let-7* microRNAs via these interactions to modulate their biological activities*. Let-7* miRNAs are known to play a role in cardiovascular diseases including cardiac hypertrophy, cardiac fibrosis, dilated cardiomyopathy and myocardial infarction (Bao *et al*, 2013).

To test the role of *H19’s let-7* binding in preventing cardiomyopathy, we used CRISPR/Cas9 genome editing to delete *let-7* binding sites in the *H19* gene (Figure 1A, 6A). Mice carrying this mutation (*H19ΔLet7 /H19+*) express *H19* at wild type levels (Figure 6B), which shows that the deletions do not disrupt lncRNA expression or stability. Adult *H19ΔLet /H19+*mice displayed cardiomegaly as measured by increased heart weight/tibia length ratios (wild type = 7.5±1.7 mg/mm, N= 4; *H19ΔLet /H19+*= 9.9±2.2 mg/mm, N=5; p = 0.007). Transverse sections also suggested hypertrophy (Figure 6C), which was quantified as increased fiber diameter (Figure 6D). The cardiac hypertrophy in *H19ΔLet7/H19+* mice was accompanied by increased interstitial and perivascular fibrosis (Figure 6E, F). The pathologic nature of the observed cardiac myopathies in these mutant mice was confirmed by the increased levels of cardiomyopathy markers (Figure 6G). Our results support a role for *let-7* miRNA binding to *H19 lncRNA* in preventing cardiomyopathies.

**Figure 6.**
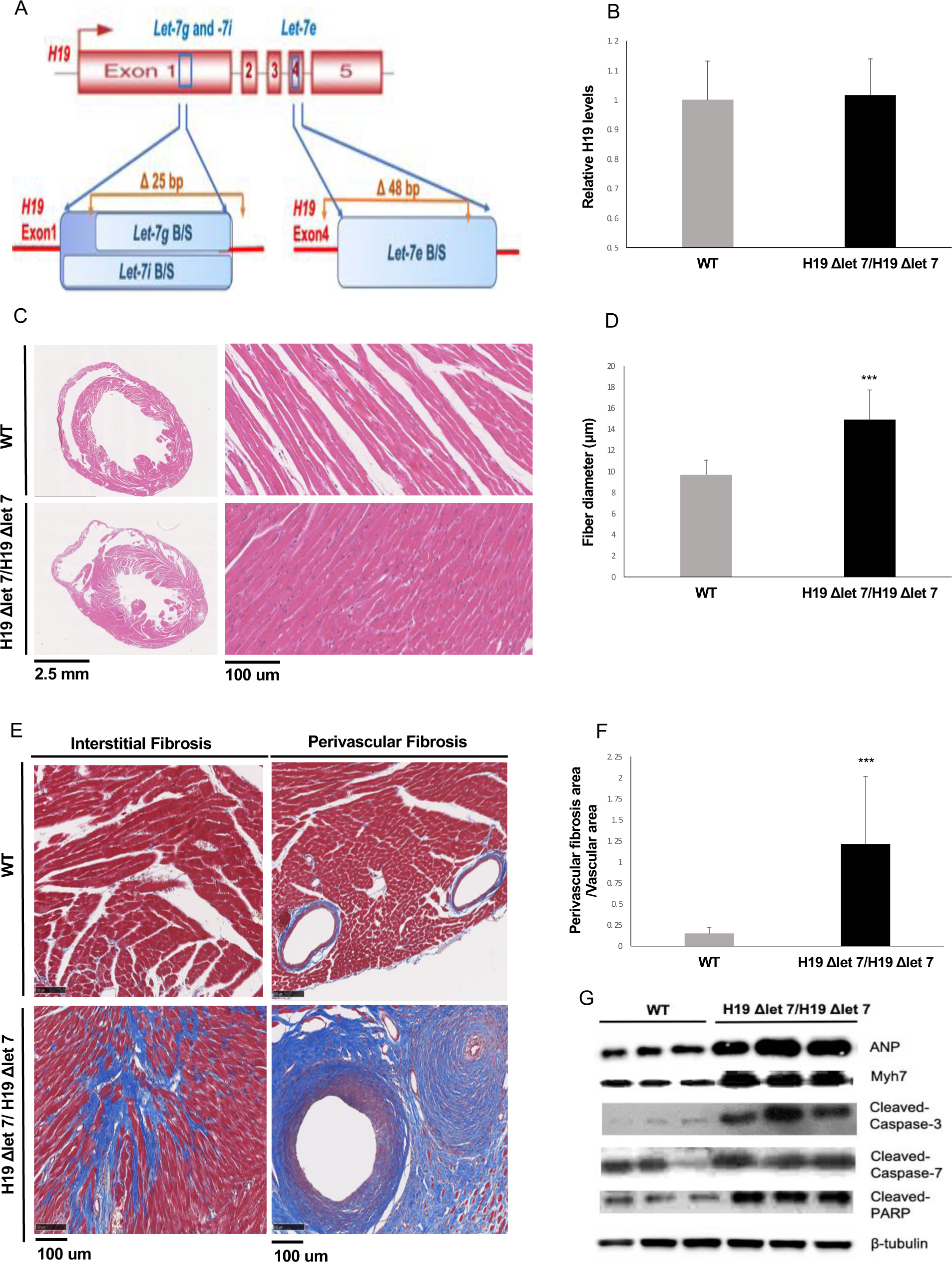
*H19*’s let7 binding domains are essential for normal function. **A.** The *H19ΔLet7* allele was generated by deleting 25 and 48 bp sequences within exons 1 and exon 4 to eliminate binding sites for *let-7g*, *let-7i,* and *let-7e* miRNAs. **B.** The *H19ΔLet7* allele is expressed at wild type levels. RNAs were isolated from hearts from *H19^ΔLet7^/H19^+^* neonates and quantitated by qRT-PCR, normalizing first to *GAPDH* and then to the levels of *H19* observed in *H19^+^/H19^+^* littermates. **C.** Transverse sections were collected midway along the longitudinal axis from hearts collected from 12-15 month old wild type (N = 4) and mutant (N = 3) littermates and stained with hematoxylin and eosin. **D.** Fiber diameters were quantitated using 3 sections per mouse. **E, F**. Masson’s trichrome staining of sections described in panel **C**. Red, muscle fibers; blue, collagen. Sections from 3 wild type and 4 mutant littermates were used to calculate fibrosis. **G.** Immunoblot analyses of whole heart extracts prepared from 12-15 month WT (N = 3) and mutant littermates (N = 3). Altered expression of ANP, Myh7, Cleaved Caspase-3, Cleaved Caspase-7, and Cleaved PARP. β-tubulin is a loading control. For all bar graphs, data are presented as mean + SEM. ***, p<0.001 (Student’s t-test).

## Discussion

BWS is an overgrowth disorder with significant patient to patient variation in disease symptoms(Jacob *et al*., 2013). An explanation for some of this variability is that independent molecular mechanisms for BWS exist(Weksberg *et al*, 2010). More than 50% of BWS cases are associated with epigenetic lesions that disrupt expression of *CDKN1C*, an imprinted gene closely linked to *IGF2/H19* but under control of its own *ICR (IC2)*. (More rarely, BWS cases are associated with pathogenic lesions in the *CDKN1C* peptide coding sequences). About 5% of BWS cases are associated with disrupted imprinting at the *IGF2/H19* locus. About 20% of cases are associated with paternal uniparental disomy of the entire region (potentially affecting both *CDKN1C* and *IGF2/H19*) (IC1 and IC2), and another 20% of cases are of unknown origin. Use of artificial reproductive technologies (ART) is a 6-10-fold risk factor for BWS specifically because of the increased chance that *IGF2/H19* imprinting is disrupted (Hattori *et al*., 2019; Johnson *et al*., 2018; Mussa *et al*., 2017). Interestingly, BWS patients associated with ART are more likely to show cardiac problems (Tenorio *et al*., 2016), suggesting a role for *IGF2/H19* expression in normal heart development and function. In this study we characterize a mouse model for *Igf2/H19* loss of imprinting (LOI). This model deletes the *Imprinting Control Region* upstream of the *H19* promoter and recapitulates the molecular phenotype of BWS patients: biallelic (i.e. 2X dosage) *IGF2* and reduced *H19* RNA. Here we show that this model phenocopies the transient cardiomegaly observed in neonates but also displays cardiovascular dysfunctions that are only rarely observed in patients.

To elucidate the molecular and developmental etiology of these cardiovascular phenotypes we characterized two additional mouse models that independently altered expression of *Igf2* and of *H19*. These genetic analyses demonstrated that overexpression of *Igf2* and loss of *H19* play distinct roles in driving BWS cardiac phenotypes. In neonates, increased levels of circulating IGF2 results in hyperactivation of mTOR signaling in cardiomyocytes and thus leads to cardiomyocyte hyperplasia and cellular hypertrophy but the resultant cardiomegaly in mice is transient. As in humans, expression of *Igf2* in mice is strongly downregulated after birth and organ sizes return toward normal. Loss of *H19*, however, results in progressive cardiac pathology. Aged *H19* deficient mice show increased fibrosis, expression of markers indicative of cardiac failure, abnormal echocardiography phenotypes, low blood pressure, and aberrant vasculature. Thus in the mouse LOI BWS model, disease phenotypes are not restricted to fetal and neonatal stages. It will be interesting and important to assess whether this is true in other mammals.

In hearts, *H19* expression is concentrated in endothelial cells. To understand the significance of *H19* expression we isolated cardiac ECs from wild type and mutant neonates. Transcriptome analyses showed altered expression of genes associated with endothelial to mesenchymal transition suggesting that *H19* might help regulate EC cell fate. Supporting this idea, we saw that forcing expression of *H19* in primary ECs prevents activation of mesenchymal gene expression patterns. Finally, we saw that *H19*-deficient mice show significant increases in the frequency of EC cells simultaneously expressing mesenchymal markers.

The ability of some ECs to transition of mesenchymal cells is necessary for normal development and thus can be assumed to be an essential property of ECs. The phenotypes of *H19*-deficient mice do <u>not</u> suggest that *H19* lncRNA is the single key molecule regulating EC cell fate: Even in LOI mice, EndMT is almost always occurring only when developmentally appropriate. Rather, our data indicate that *H19* RNA levels play a role in modulating the fate decision so that cells lacking *H19* are modestly but measurably more likely to switch toward a transitional state where both EC and mesenchymal markers are expressed. It is interesting to note one commonality of key pathways disrupted by loss of *H19* lncRNA is that they share regulation by TGF*β* signaling, suggesting that the observed 50% reduction in expression of TGF*β* receptors might be a key phenotype in *H19*-deficient ECs (Goumans *et al*., 2008).

Our findings extend earlier studies showing patterns of *H19* expression in development and in response to injury suggesting a role for *H19* in vascular physiology and pathology (Jiang *et al*, 2016; Kim *et al*, 1994). Moreover, Voellenkle et al. recently described a role for lncRNAs including *H19* in the physiology of umbilical vein endothelial cell (Voellenkle *et al*, 2016).

Our results also agree with in vitro studies that demonstrated an important role for *H19* in regulating EMT in cancer cells (Li *et al*, 2019; Ma *et al*, 2014; Matouk *et al*, 2016; Matouk *et al*, 2014; Wu *et al*, 2019; Zhang *et al*, 2018) In these previous analyses, *H19* function was determined by transfecting cancer cells with *H19*-expression vectors and analyzing cell motility and gene expression. However, in contrast to our findings that activation of mesenchymal expression is associated with <u>loss</u> of *H19*, these in vitro analyses find EMT is induced by <u>increasing</u> *H19* RNA. This discrepancy emphasizes the useful role for genetic animal models in addressing developmental disorders where phenotypes are coming from cumulative changes in multiple cell types and over long periods of time.

In animal models, observed phenotypes are due to the cumulative effect of the mutation in many cell types and over developmental time. In vitro studies of H19 have focused on the effect of acute changes in levels of *H19* in a single cell type. Th Our analyses cannot address the fate of these cells that co-express EC and mesenchymal markers. Do they all proceed toward full EndMT, do they return toward EC fates, or do they teeter in between? These questions can be addressed in future experiments using conditional *H19* deletion alleles and cell fate markers.

*H19* can be a very abundant transcript. In neonatal ECs, *H19* lncRNA represents about 1% of all polyadenylated RNA. Yet its biochemical functions remain unclear. Various studies support the idea that *H19* functions as a microRNA precursor (Cai & Cullen, 2007; Dey *et al*, 2014; Keniry *et al*., 2012), a p53 protein inhibitor (Hadji *et al*., 2016; Park *et al*., 2017; Yang *et al*., 2012; Zhang *et al*., 2017), a regulator of DNA methylation (Zhou *et al*., 2019; Zhou *et al*., 2015), and as a modulator of *let7* microRNA functions (Gao *et al*., 2014; Geng *et al*., 2018; Kallen *et al*., 2013; Peng *et al*., 2017; Zhang *et al*., 2019; Zhang *et al*., 2017). It is possible that *H19* functions vary from cell type to cell type (Raveh *et al*, 2015). Alternatively, these functions might co-exist in a single cell but analyses to date have only looked at *H19* function from single perspectives and have missed its ability to perform in multiple pathways. To address this issue, we have begun to generate mutant *H19* alleles that disrupt specific functions. Here we show that *H19ΔEx1/H19^+^* mice have 100X reduced levels of lncRNA but almost normal levels of *mi675* and still show cardiac pathology. Thus, the pathologies in LOI mice depend on the loss of *H19* lncRNA. To then address how the lncRNA might function, we generated mice carrying an *H19* allele missing *let7* binding sites. These mice show cardiac pathologies including extreme fibrosis. We find the fibrosis phenotype in *H19ΔLet7* mice to be especially interesting. We speculate that intensity relative to that seen in an *H19* null is most consistent with the idea that *H19* lncRNA has multiple roles in the cell and by disrupting only one role we have altered some balance so that the animal is worse off than having no *H19* at all.

The strong phenotype in *H19ΔLet7* mice is consistent with several previous studies that emphasize the importance of *H19* lncRNA interactions with *let-7* miRNAs but it is also paradoxical in that multiple studies of let7 function in hearts indicate that *let-7* functions as an anti-fibrotic factor. That is, reduced *let-7* is a risk factor for fibrosis and fibrosis induces *let-7*, presumably as a corrective measure (Bao *et al*., 2013; Elliot *et al*, 2019; Sun *et al*, 2019; Wang *et al*, 2015). The increased fibrosis in *H19ΔLet7*-mice suggests that the simple model (that *H19* binds to and reduces *let-7* bioavailability) is not correct or, more likely, that complex developmental interactions play critical roles that determine phenotypes in ways that are not yet understood. Either way, our results confirm the importance of animal models and the need for even more sophisticated conditional deletions.

*H19* and *Igf2* are generally thought of as fetal genes since their expression is so strongly repressed after birth. This fact might suggest that the adult phenotypes in *H19*-deficient mice are downstream effects of the loss of *H19* in the developing heart. However, as already mentioned, at peak expression, *H19* levels are extraordinarily high. Thus, even after 100-fold developmentally regulated decrease, *H19* remains one of the top 100 genes in terms of RNA levels. For this reason, conditional ablation models will be needed to determine exactly when *H19* expression is important.

## Materials and Methods

### Animal Studies

All mice were bred and housed in accordance with National Institutes of Health and United States Public Health Service policies. Animal research was performed only after protocols were approved by the National Institute of Child Health and Human Development Animal Care and Use Committee.

*H19^ΔICR^/H19^+^* (Srivastava *et al*., 2000) and wild type littermates or *H19^ΔEx1^/H19^+^* (Srivastava *et al*., 2003) and wild type littermates were generated by backcrossing heterozygous females with C57BL/6J males (Jackson Labs 000664). For tissue specific LOI, we crossed *H19^ΔICR^/H19^ICRflox^* females (Srivastava *et al*., 2000) with males hemizygous for the *Myh6Cre* transgene (Jackson Labs 011038) (Agah *et al*, 1997). The H19 BAC transgene was generated as described (Kaffer *et al*., 2001; Kaffer *et al*., 2000) and used to generate *H19^ΔICR^/H19^+^ BAC+* females for backcrosses with C57BL/6J males.

The *H19ΔLet7* allele was generated using CRISPR/Cas9 gene editing of RI mouse embryonic stem cells (ESCs). In step 1, we used gRNAs 5’-CACCGAGGGTTGCCAGTAAAGACTG-3’ and 5’-CACCGCTGCCTCCAGGGAGGTGAT - 3’to delete 25 bp (AGACTGAGGCCGCTGCCTCCAGGGAGGTGAT) in exon 1. In step 2, we used gRNAs: 5’-CACCGCTTCTTGATTCAGAACGAGA-3’ and 5’-CACCGACCACTACACTACCTGCCTC-3’ to delete 48 bp (CGTTCTGAATCAAGAAGATGCTGCAATCAGAACCACTACACTACCTGC) in exon 4. Positive clones were identified by PCR screens and then confirmed by sequencing 686 bp spanning the exon 1 deletion and 1086 bp spanning the exon 4 deletion. Founder mice were obtained by injecting mutated ESCs into C57BL/6J blastocyts and then backcrossed twice to C57BL/6J females.

Genotypes were determined by PCR analyses of gDNAs extracted from ear punch biopsies (Supplemental Table 2).

### Electrocardiography measurements

Transthoracic echocardiography was performed using a high-frequency linear array ultrasound system (Vevo 2100, VisualSonics) and the MS-400 Transducer (VisualSonics) with a center operating frequency of 30 MHz, broadband frequency of 18 to 38 MHz, axial resolution of 50 mm, and footprint of 20×5 mm. M-mode images of the left ventricle were collected from the parasternal short-axis view at the midpapillary muscles at a 90° clockwise rotation of the imaging probe from the parasternal long-axis view. Form the M-mode images, the left ventricle systolic and diastolic posterior and anterior wall thicknesses and end-systolic and -diastolic internal left ventricle chamber dimensions were measured using the leading-edge method. Left ventricular functional values of fractional shortening and ejection fraction were calculated from the wall thicknesses and chamber dimension measurements using system software. Mice were imaged in the supine position while placed on heated platform after light anesthesia using isoflurane delivered by nose cone.

### Blood Pressure measurements

After sedation with isoflurane, a pressure catheter (1.0-Fr, model SPR1000, Millar Instruments, Houston, TX) was inserted into the right carotid and advanced to the ascending aorta. After 5-minute acclimation, pressures were recorded using Chart 5 software (AD Instruments, Colorado Springs, CO) (Knutsen *et al*, 2018).

### Arterial Pressure-Diameter Testing

Ascending aortas (from the root to just distal to the innominate branch point) and left carotid arteries (from the transverse aorta to 6 mm up the common carotid) were dissected and mounted on a pressure arteriograph (Danish Myotechnology, Copenhagen, Denmark) in balanced physiological saline (130 mM NaCl, 4.7 mM KCl, 1.6 mM CaCl_2_, 1.18 mM MgSO_4_·7H_2_O, 1.17 mM KH_2_PO_4_, 14.8 mM NaHCO_3_, 5.5 mM dextrose, and 0.026 mM EDTA, pH 7.4) at 37°C. Vessels were transilluminated under a microscope connected to a charge-coupled device camera and computerized measurement system (Myoview, Danish Myotechnology) to allow continuous recording of vessel diameters. Prior to data capture vessels were pressurized and stretched to in vivo length(Wagenseil *et al*, 2005). Intravascular pressure was increased from 0 to 175 mmHg in 25-mmHg steps. At each step, the outer diameter (OD) of the vessel was measured and manually recorded. Segmental distensibility was calculated from the pressure diameter curves as follows: distensibility (SD_25_) over a 25-mmHg interval = [OD_Higher Pressure (H)_ − OD_Lower Pressure(L)_]/OD_(L)_/25. (Knutsen *et al*., 2018).

### Histological Analyses

Hearts from adult mice were fixed by Langendorff perfusion or by transcardiac perfusion using 4% paraformaldehyde ((PFA) and embedded in paraffin. Fetal and neonatal hearts were isolated and then fixed by submersion in 4% PFA before embedding. From embedded hearts, we obtained 5 mm transverse sections for analysis. Masson’s Trichrome (Sigma Aldrich, HT15, St. Louis Missouri) and Picosirius Red (Sigma Aldrich, 365548) staining were according to supplier’s instructions. Fiber diameter index was quantitated using Hamamatsu-NDP software. *Immunofluorescence and Immunohistochemistry* Primary myocytes and H19C2 cells were fixed with 4% PFA, permeabilized with 0.5% Triton, and blocked with 10% normal serum before incubation with antibodies. Paraffin sections were deparaffinized and rehydrated according to standard protocols. Antigen retrieval was applied using citrate buffer (Abcam, 1b93679, Cambridge, MA) for 20 minutes and then maintained at a sub-boiling temperature for 10 minutes. Sections were treated with serum-free blocking solution (DAKO, X0909, Santa Clara, CA) and all antibodies (Supplemental Table 3) diluted in antibody diluent solution (DAKO, S0809). Secondary staining was performed for 30 min. at RT. Samples were imaged with a Carl Zeiss 880 laser scanning microscope using a 40X oil immersion objective. Images were composed and edited in ZEN&LSM image software provided by Carl Zeiss or Illustrator 6.0 (Adobe).

### RNA in situ hybridization

Single color probes for H19 were purchased from Advanced Cell Diagnostics (ACD 423751, Newark, CA). RNA in-situ hybridization was performed on paraffin sections using the 2.5 HD Brown Detection Kit (ACD 322310). For dual staining with antibodies, we used H19-RD chromagen kit (ACD 322360).

### Immunoblotting

Cell extracts and tissue extracts were prepared using M-PER mammalian protein extraction buffer (Thermo Fisher 78501, Waltham, MA) or T-PER tissue protein extraction buffer (Thermo Fisher 78510), respectively. Protein concentrations were assayed using a BCA Protein Assay Kit (Pierce 23227, Waltham, MA). Proteins were fractionated by electrophoresis on 12% or on 4-20% SDS-PAGE gels and then transferred to nitrocellulose. Antibodies (Supplemental Table 3) were diluted in antibody enhancer buffer (Pierce 46644).

### Cell culture

Primary cardiomyocytes were isolated form P1 pups using the Pierce Primary Cardiomyocyte Isolation Kit (Thermo Fisher 88281). H19c2 cells were purchased from ATCC (CXRL-1446). and grown at 37°C in 5% CO_2_ in DMEM + 10% FBS. Cell surface index was quantitated using Carl Zeiss-LSM software (n = 50 for each of 3 independent experiments).

To prepare primary endothelial cells, neonatal hearts were isolated and dissociated into single cells using Miltenyi Biotec Neonatal Heart Dissociation Kit (130-098-373, Gaithersburg, MD) but omitting the Red Cell Lysis step. Endothelial cells were purified based on CD31 expression (Miltenyi Biotec Neonatal Cardiac Endothelial Cell Isolation Kit, 130-104-183).

### Quantitative real-time PCR for RNA samples

Conventional RNAs were prepared from 3-5 independent biological samples, analyzed using a Thermo Fisher NANODROP 2000c to evaluate purity and yield, and then stored at −70°C. cDNA samples were prepared with and without reverse transcriptase using oligo-dT primers (Roche, 04 887 352 001). cDNAs were analyzed using SYBR Green (Roche, 04 887 352 001) on the Roche Light Cycler 480 II (45 cycles with annealing at 60C) using primers described in Supplemental Table 2. For each primer pair, we established standard curves to evaluate slope, y-intercepts, and PCR efficiency and to determine the dynamic range of the assay. Assay specificity was determined by melting point analyses and gel electrophoresis.

For microRNA analyses we used mirVanaTM miRNA Isolation Kit and TaqMan MicroRNA Assays (Thermo Fisher, 4437975; Assay ID 001973 (U6), 001940 (miR-675-5p), 001941 (miR-675-3p).

### ELISA

IGF2 secreted peptide was assayed with the Mouse IGF2 ELISA KIT (Abcam, ab100696) on 10 independent samples.

### RNA sequencing and analyses

For analyses in adult animals, RNAs were isolated from 6-month H19*Δ*ICR/H19*Δ*ICR and H19+/H19+ littermates (2 per genotype) using RNeasy Plus Mini Kit (Qiagen). Samples with RNA Integrity numbers >9 were Ribosomal RNA depleted using RiboZero Gold Kit (Illumina). Libraries were prepared using an RNA Sample Prep V2 Kit (Illumina), were sequenced (Illumina HiSeq2500) to generate paired-end 101 bp reads that were aligned to the mouse genome version mm10 using STAR v2.5.3a (Dobin et al 2013). Differential expression analyses were performed using DESeq2 (Love *et al*, 2014).

For analyses in neonates, RNAs were isolated from purified cardiac endothelial cells isolated from *H19^ΔEx1^/H19^+^* (N=3) and *H19^+^/H19^+^* (N=4) littermates. Libraries were generated from samples with RNA Integrity Numbers >9 and were sequenced and analyzed as described above.

## Acknowledgements

We thank Victoria Biggs and Jeanne Yimdjo for animal husbandry. This work was supported by the Division of Intramural Research of the Eunice Kennedy Shriver National Institute of Child Health and Human Development.

## Author Contributions

Ki-Sun Park, Beenish Rahat, Russell Knutsen, and Karl Pfeifer designed and performed experiments, analyzed data, and wrote the manuscript. Zu-Xi Yu and Danielle Springer designed and performed experiments and analyzed data. Jacob Noeker analyzed data and wrote the manuscript. Apratim Mitra analyzed data. Beth Kozel analyzed data and wrote the manuscript.

## Data Availability

RNA sequencing data are deposited in the NCBI Gene Expression Omnibus (GEO) under Series Accession number GSE111418.

---

**Supplemental Figure 1.**
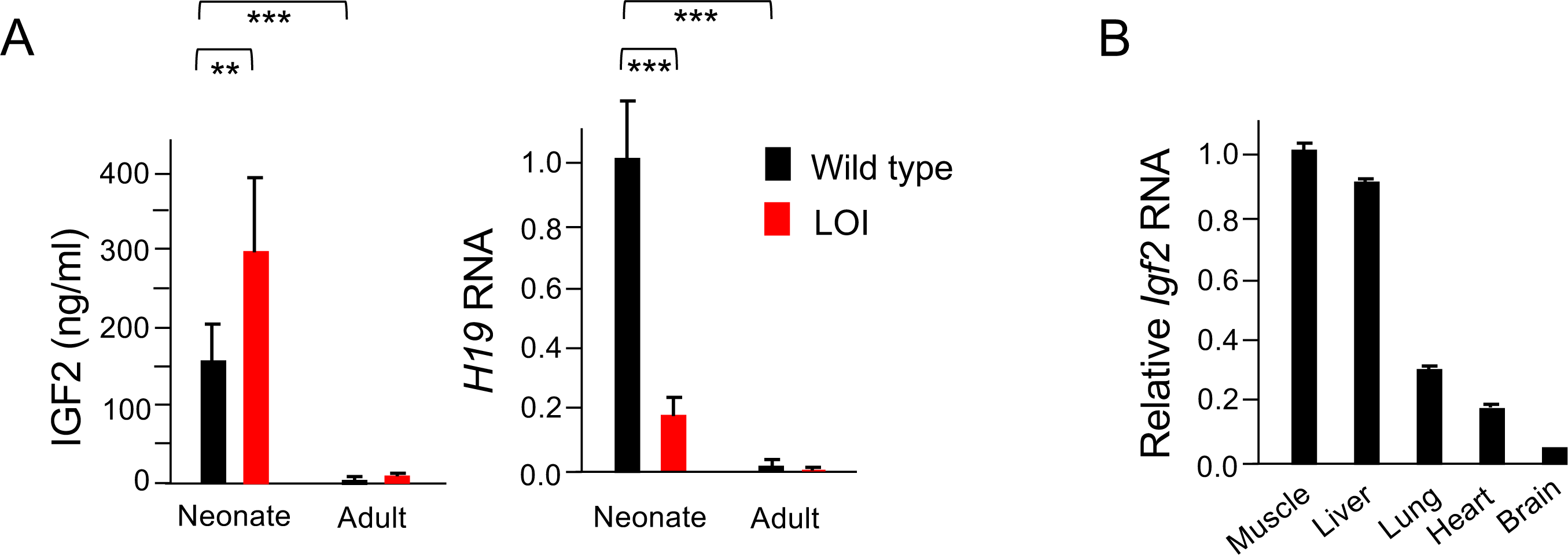
I*g*f2 and *H19* expression in wild type and LOI mice. A Maternal Loss of Imprinting (LOI) results in 2X IGF2 and reduced *H19* lncRNA. IGF2 peptide levels in serum were measured by ELISA (N=10). To quantitate *H19*, RNAs were extracted from total hearts, analyzed by qRT-PCR, normalized to GAPDH, and then normalized to RNA levels observed in wild type neonates (N > 4). Despite the dramatic postnatal repression, *H19* expression in adults remains substantial and *H19* RNA is among the top 10-percentile of all RNAs. B *Igf2* levels vary by tissue. RNAs were extracted from hind limb muscle, liver, lung, whole heart, and brain from P2 neonates and quantitated as above but normalized to *Igf2* levels in hind limb muscle. A, B Data are presented as mean + SEM. **, p<0.01; ***, p<0.001 (Student’s t-test).

**Supplemental Figure 2.**
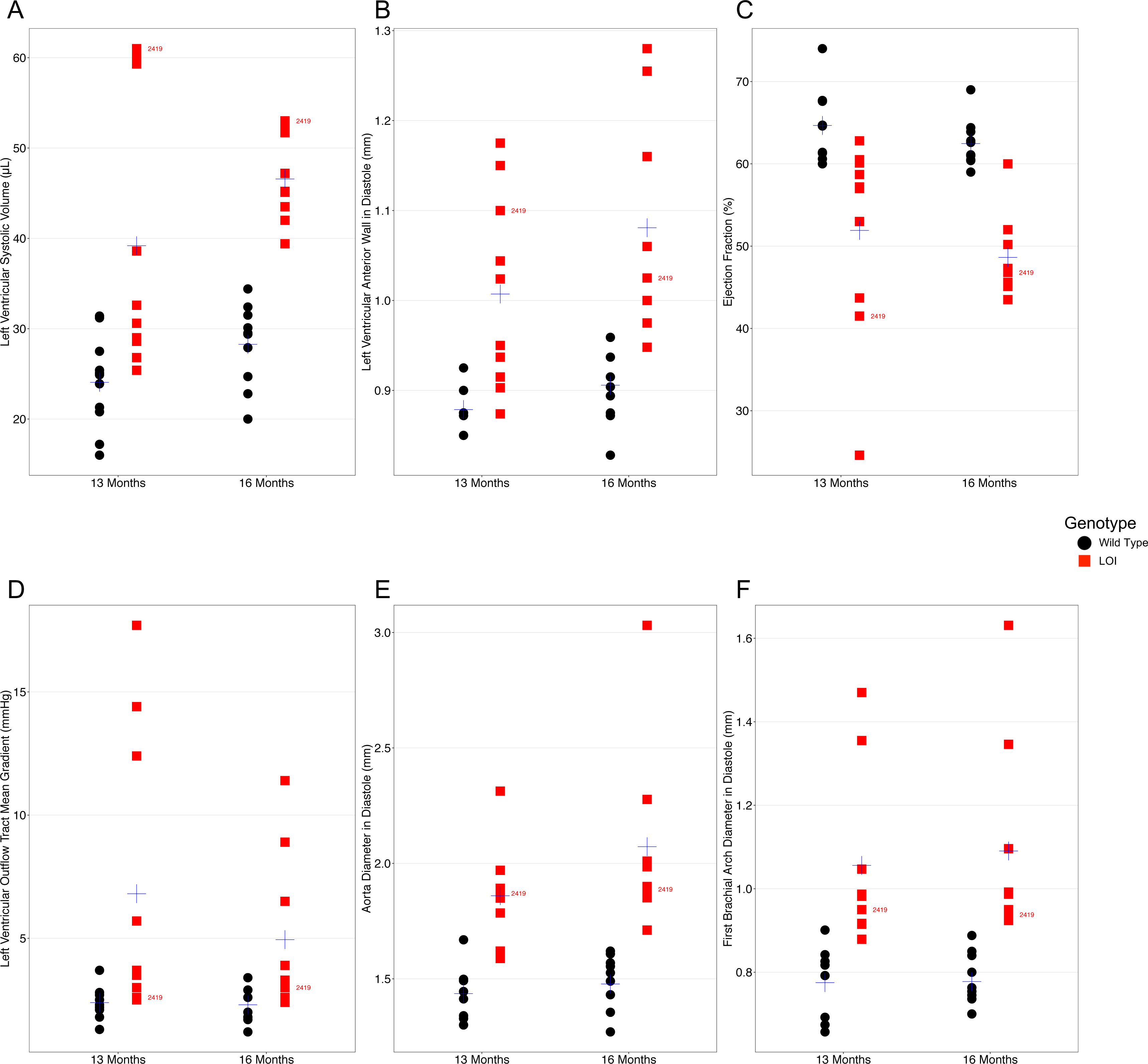
Echocardiography measures from 11 wild type and 10 LOI mice at 13 months and for 11 wild type and 9 LOI mice at 16 months. Loss of imprinting results in independent functional and vascular phenotypes. A, B, C Left ventricular phenotypes are progressive and stratify with time. D, E, F Arterial defects, abnormal outflow and increased artery diameters, do not progress. Arterial diameter phenotypes are already stratified at 13 months. A-F Black dots represent wild type and red squares represent LOI mice. Blue crosses represent mean. One sample (mouse 2419) is labelled to show that extreme phenotypes do not correlate. B *Igf2* levels vary by tissue. RNAs were extracted from hind limb muscle, liver, lung, whole heart, and brain from P2 neonates and quantitated as above but normalized to *Igf2* levels in hind limb muscle. A, B Data are presented as mean + SEM. **, p<0.01; ***, p<0.001 (Student’s t-test).

**Supplemental Figure 3.**
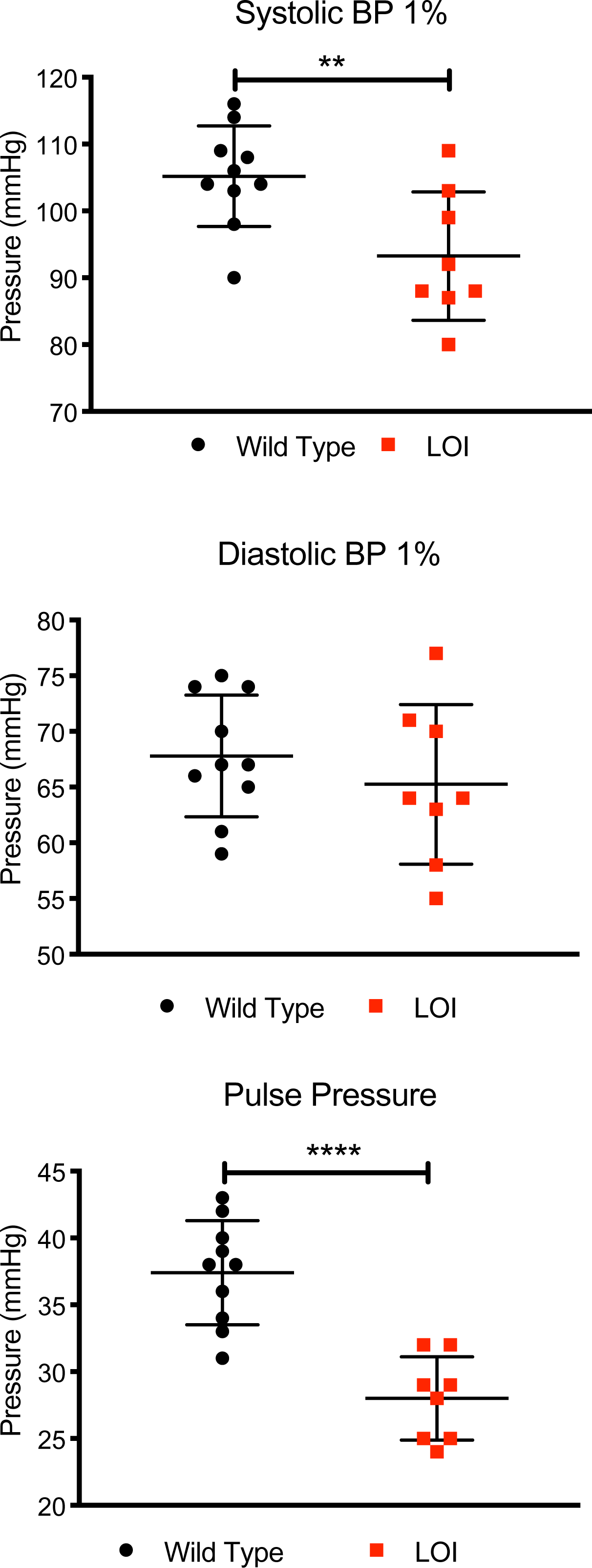
Decreased systolic and pulse pressures in LOI mice. Blood pressures from 10 wild type and 8 LOI sedated mice were measured as described in Methods. Means and standard deviation are shown as whiskers underlying the data points. Systolic and diastolic blood pressure data were analyzed by Tukey’s test; pulse pressure data were analyzed using one-way ANOVA. **, P < 0.01; ****, P < 0.0001. Diastolic pressures were not significantly different. A-F Black dots represent wild type and red squares represent LOI mice. Blue crosses represent mean. One sample (mouse 2419) is labelled to show that extreme phenotypes do not correlate. B *Igf2* levels vary by tissue. RNAs were extracted from hind limb muscle, liver, lung, whole heart, and brain from P2 neonates and quantitated as above but normalized to *Igf2* levels in hind limb muscle. A, B Data are presented as mean + SEM. **, p<0.01; ***, p<0.001 (Student’s t-test).

**Supplemental Figure 4.**
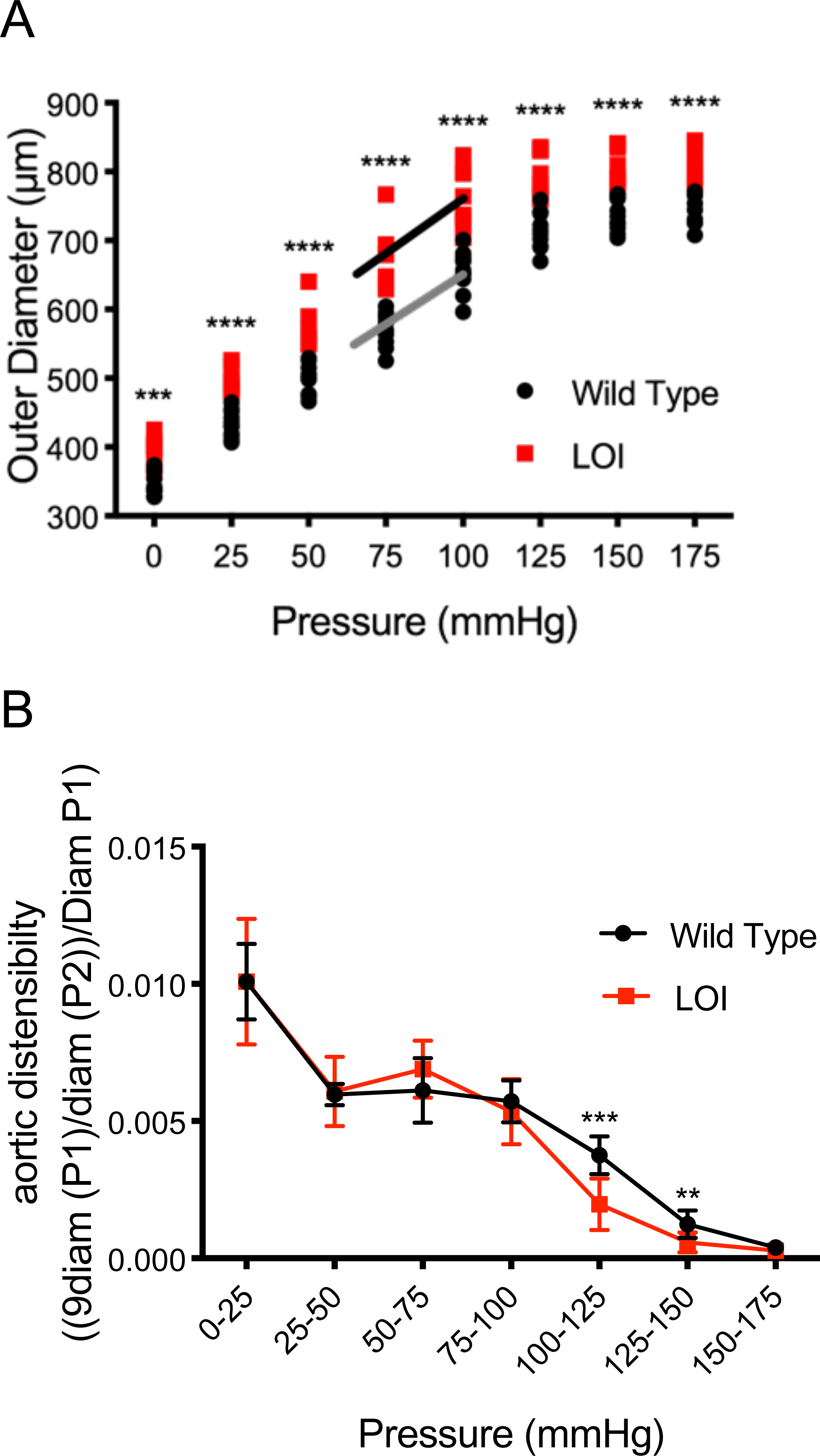
Increased vessel diameter and distensibility in carotid arteries isolated from 10 wild type and 8 LOI mice at 16-months. A Increased diameters across a wide range of applied pressures. B Increased segmental distensibility across physiologically relevant pressures. A, B **, P<0.01; ***, P<0.001; ****, P<0.0001 (Two-way repeated measure ANOVA). B *Igf2* levels vary by tissue. RNAs were extracted from hind limb muscle, liver, lung, whole heart, and brain from P2 neonates and quantitated as above but normalized to *Igf2* levels in hind limb muscle. A, B Data are presented as mean + SEM. **, p<0.01; ***, p<0.001 (Student’s t-test).

**Supplemental Figure 5.**
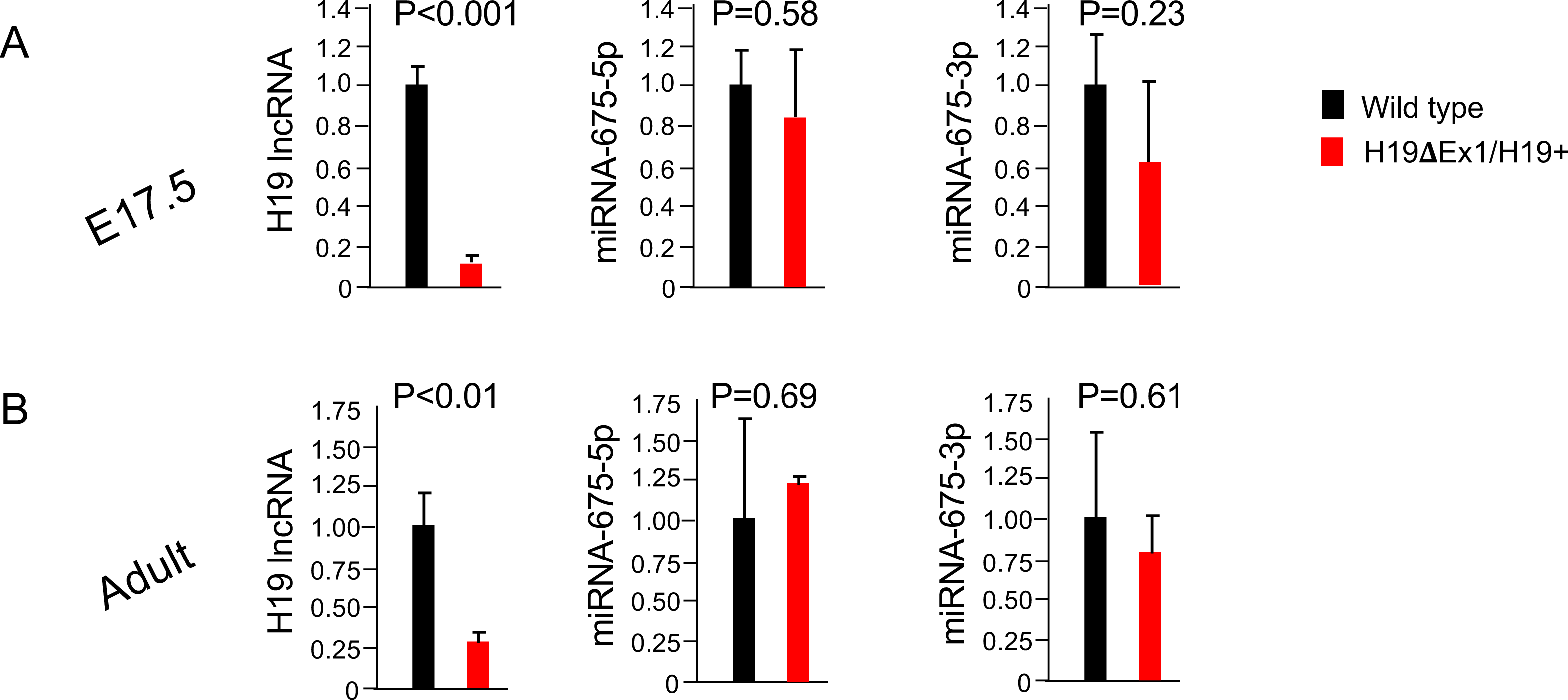
The H19ΔEx1 deletion specifically reduces H19 lncRNA. RNAs were extracted from whole hearts isolated from wild type and from *H19^ΔEx1^/H19^+^* littermates at e17.5 and at 1 year of age. *H19* lncRNA levels were normalized to GAPDH. mi675 levels were normalized to U6 miRNA. In each experiment, expression in *H19^ΔEx1^/H19^+^* is normalized to the wild type littermates. Data are presented as mean + SEM (Student’s t-test). N > 3. B *Igf2* levels vary by tissue. RNAs were extracted from hind limb muscle, liver, lung, whole heart, and brain from P2 neonates and quantitated as above but normalized to *Igf2* levels in hind limb muscle. A, B Data are presented as mean + SEM. **, p<0.01; ***, p<0.001 (Student’s t-test).

**Supplemental Figure 6.**
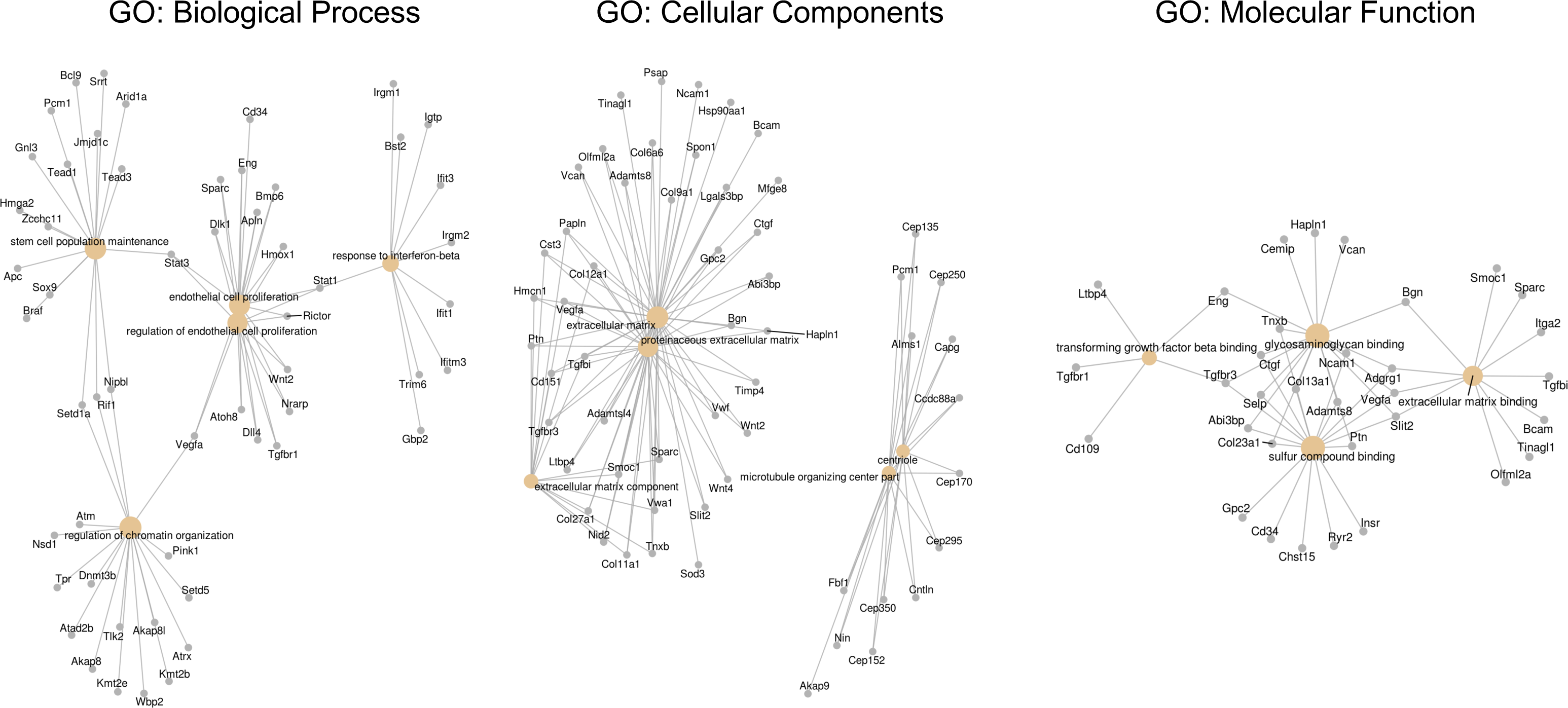
Visual representation of Gene Ontology analysis for endothelial cells isolated from wild type (N = 4) and *H19ΔEx1/H19+* (N = 3) P2 neonatal hearts. RNAs were isolated and polyadenylated transcripts were quantitated. Cnetplot depicts linkages of genes and biological concepts as a network. Nodes identify complex associations of genes that contribute to a functional term within a pathway. GO: Biological Process indicates enriched mesenchymal biological markers and regulation of endothelial proliferation genes. GO: Cellular Components indicate enrichment in extracellular matrix genes. GO: Molecular function indicates enrichment in extracellular matrix and TGF-beta binding genes.

**Supplemental Table 1.** Correlations between echocardiography phenotypes in 13 month old mice. As displayed in Supplemental Figure 2, each mouse presents a unique array of phenotypes. Increased diameter sizes for major vessels is.

|  | Ejection Fraction (%) | LV Volume (sys) | LVAW (dia) | LVOT mean velocity | Aorta diameter (sys) | Brachial Arch diameter (syst) |
| --- | --- | --- | --- | --- | --- | --- |
| Ejection Fraction (%) | 1.000 | 0.810 | 0.526 | 0.024 | 0.005 | 0.155 |
| LV Volume (sys) |  | 1.000 | 0.495 | 0.002 | 0.097 | 0.158 |
| LVAW (dia) |  |  | 1.000 | 0.287 | 0.068 | 0.441 |
| LVOT mean velocity |  |  |  | 1.000 | 0.430 | 0.623 |
| Aorta diameter (sys) |  |  |  |  | 1.000 | 0.680 |
| Brachial Arch diameter (sys) |  |  |  |  |  | 1.000 |

**Supplemental Table 2.**
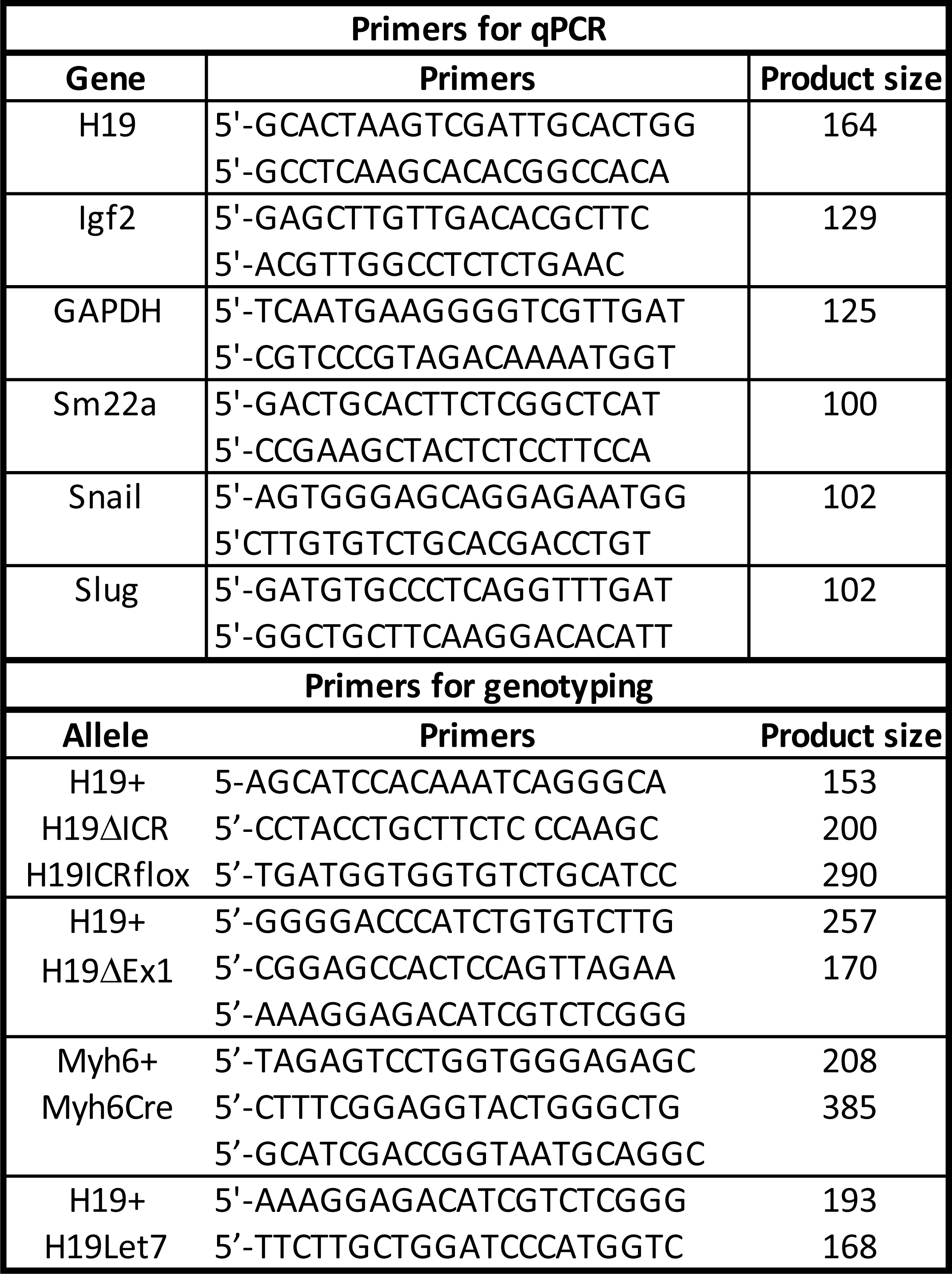
Primers used for qRT-PCR and for genotyping.

**Supplemental Table 3.** Antibodies used in this study.

| Antibody | Company | Catalog # | Dilution |
| --- | --- | --- | --- |
| ANP | Abcam | ab91250 | 1:1000 |
| Myh7 | Santa Cruz | sc52089 | 1:200 |
| Serca2 | Abcam | ab2861 | 1:1000 |
| Cleaved Caspase 3 | Abcam | ab9664 | 1:1000 |
| Cleaved Caspase 7 | Abcam | ab8438 | 1:1000 |
| Cleaved Parp | Abcam | ab5625 | 1:1000 |
| β-tubulin | Abcam | ab6046 | 1:2000 |
| Ki-67 | Abcam | ab15580 | 1:1000 |
| Cyclin E1 | Abcam | ab3927 | 1:100 |
| Cyclin D1 | Abcam | ab134175 | 1:1000 |
| GAPDH | Cell Signalling | 2118 | 1:2000 |
| Myh6 | Santa Cruz | sc168676 | 1:100 |
| Phalloidin | Cell Signalling | 8593 | 1:200 |
| p-AKT | Cell Signalling | 4060 | 1:1000 |
| t-AKT | Cell Signalling | 9272 | 1:1000 |
| p-S6K1 | Cell Signalling | 9234 | 1:1000 |
| t-S6K1 | Cell Signalling | 2708 | 1:1000 |
| p-rpS6 | Cell Signalling | 4858 | 1:1000 |
| t-rpS6 | Cell Signalling | 2317 | 1:1000 |
| CD31 | Abcam | ab28364 | 1:50 |
| aSMA | Abcam | 1b21027 | 1:100 |
| aSMA | GeneTex | GTX25694 | 1:500 |
| vWF | DAKO | A0082 | 1:1000 |
| 488 Goat anti-rabbit | Life Technology | A11008 | 1:500 |
| 488 Goat anti-mouse | Life Technology | A11001 | 1:500 |
| 488 Donkey anti-goat | Life Technology | A11055 | 1:500 |
| <u>555 Goat anti-mouse</u> | <u>Life Technology</u> | <u>A21424</u> | <u>1:500</u> |
| <u>555 Donkey anti-Rabbit</u> | <u>Life Technology</u> | <u>A31572</u> | <u>1:500</u> |
| DAPI | Life Technology | D3571 | 1:1000 |

